# Disruption of Plasma Membrane Lipid Asymmetry Alters Cellular Energetics

**DOI:** 10.1101/2025.04.27.650841

**Authors:** Pavel Barahtjan, Katelyn C. Cook, Václav Bočan, H. Mathilda Lennartz, Tobias Jumel, Kai Schuhmann, Tyler Robert Reagle, Silvia Radrezza, Sofia Traikov, Kristin Böhlig, Sascha M. Kuhn, Cristina Jiménez López, Tiago C. Alves, Andrej Shevchenko, Jonathan Rodenfels, André Nadler

**Affiliations:** Max-Planck-Institute of Molecular Cell Biology and Genetics, Dresden, 01307, Germany; Interfaculty Institute of Bioengineering and Global Health Institute, École Polytechnique Fédérale de Lausanne (EPFL), Lausanne, 1015, Switzerland; Molecular Physiology and Biological Physics, Center for Membrane and Cell Physiology, School of Medicine, University of Virginia, Charlottesville, VA 22903, USA; Institute for Clinical Chemistry and Laboratory Medicine, Universitätsklinikum Carl Gustav Carus, Technical University Dresden, Germany; DZD-Paul Langerhans Institute of the Helmholtz Zentrum München, Germany; Cluster of Excellence Physics of Life, Technical University Dresden, Dresden, Germany

## Abstract

Biological membranes often feature unequal and actively maintained concentration gradients of lipids between bilayer leaflets, termed lipid asymmetry.^1–3^ Lipid asymmetry has been linked to many cellular processes^4,5^, but the mechanisms that connect lipid trans-bilayer concentration gradients to cellular functions are poorly understood. Here we systematically map the cellular processes affected by dysregulation of lipid asymmetry by knocking out flippases, floppases, and scramblases in mammalian cells. We identify broad alterations to core metabolic pathways, including nutrient uptake, neutral lipid turnover, the pentose phosphate pathway, and glycolysis. Lipidomics, respirometry, lipid and metabolic imaging, and live-cell calorimetry revealed elevated neutral lipid and ATP consumption rates, coupled with increased heat loss per unit biomass synthesized despite slower growth. This suggests that compensatory maintenance of lipid asymmetry strains the cellular energy budget, shifting the cell from a growth-promoting anabolic state to a more energy-using catabolic state. Our data indicate that lipid asymmetry and its active maintenance are linked to cellular energetics, reflecting either the energetic cost of maintaining lipid asymmetry or its putative role as a cellular store of potential energy akin to proton and ion transmembrane gradients.

## Main Text

The plasma membranes of eukaryotes and bacteria display an asymmetric lipid distribution between the inner and outer bilayer leaflets, far from thermodynamic equilibrium^1–4,6–8^. Lipid asymmetry constitutes a trans-bilayer lipid concentration gradient. Maintaining this steady-state lipid concentration gradient requires actively driven lipid flows across the membrane, generated and controlled by lipid translocases. Specifically, flippases and floppases (P_4_-ATPases and ABC transporters) move lipids against the gradient in an ATP-dependent, substrate-specific manner.^5,9,10^ ATP-independent scramblases equilibrate lipid distribution between the bilayer leaflets (Fig. 1A).^5,11–13^

**Fig. 1.**
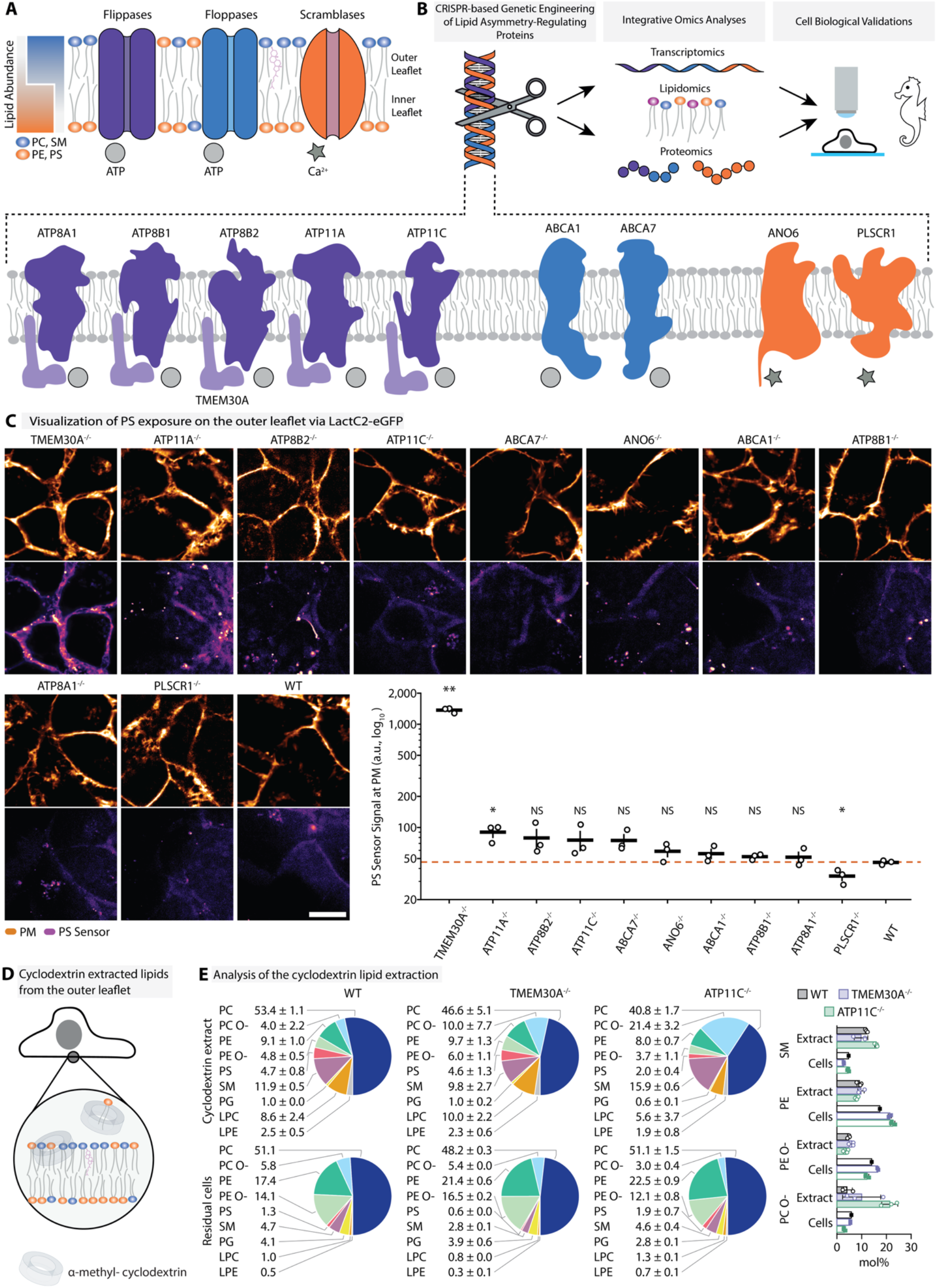
Knockout of lipid translocases alters plasma membrane lipid transbilayer distribution. **(A)** Schematic visualization of lipid asymmetry and its protein machinery. The asymmetric distribution of lipids is generated and maintained by ATP-hydrolyzing P_4_-ATPases (flippases), ABC transporters (floppases), and dissipated by Ca^2+^-dependent scramblases. These proteins drive the characteristic distribution of phosphatidylcholine (PC) and sphingomyelin (SM) to the outer leaflet and phosphatidylethanolamine (PE) and phosphatidylserine (PS) to the inner leaflet of the plasma membrane. **(B)** Experimental workflow used in this study. Lipid translocases are targeted for CRISPR-mediated gene knockouts. The generated KO cell lines are subjected to bulk transcriptomic, proteomic, and lipidomic analyses. Detected changes are further validated by biochemical and cell biological experiments. **(C)** Visualization of PS exposure on the outer leaflet of the plasma membrane of living WT and KO cells with the PS biosensor LactC2-eGFP and quantification of the LactC2-eGFP signal at the plasma membrane (bottom right). The plasma membrane was counterstained with CellMask Deep Red (PM, upper image rows) and LactC2-eGFP was applied extracellularly (PS Sensor, lower image rows). All images were acquired with the same microscopy settings. LactC2-eGFP is shown on an intensity scale calibrated on TMEM30A^-/-^ cells for TMEM30A^-/-^ and on a shared, approximately twenty times more sensitive scale for all other cell lines. Scale bar = 10 µm. Points represent individual fields (n = 3 fields per cell line from one well), horizontal lines indicate the mean and error bars the standard error of the mean. The dashed line indicates the WT mean. p-Values were calculated using a two-sided Welch t-test for each KO compared to WT and are shown without correction for multiple comparisons: ** < 0.01, * < 0.05, NS = not significant. After Benjamini-Hochberg correction for ten comparisons, only TMEM30A^-/-^ differs significantly from WT. **(D)** Extraction of lipids from the outer leaflet of the plasma membrane with methyl-α-cyclodextrin. Cartoon schematic of the extraction. **(E)** Analysis of the cyclodextrin lipid extraction. Phospholipid class composition of the cyclodextrin extracts (top) and the residual cells (bottom) of WT, TMEM30A^-/-^ and ATP11C^-/-^ cells, shown as mol% of the nine measured phospholipid classes (all species of a class summed) within each sample; labels give the mean ± SD for each class (mean only for the single WT residual-cell sample). The bar graph (right) shows SM, PE and the ether lipids PE O- and PC O-in the extracts (tinted bars, open circles) and the residual cells (white bars, filled squares). Points represent individual samples; bars indicate the mean, and error bars indicate the standard deviation; no statistical tests were applied. Species with an odd number of carbon atoms and extract species not detected in the residual cells were excluded; PS in the residual cells is likely underestimated (see Methods). Extracts: n = 3 samples for each cell line; residual cells: n = 1, 2 and 3 samples for WT, TMEM30A^-/-^ and ATP11C^-/-^, respectively.

In mammals, lipid asymmetry has been linked to numerous cellular processes, including apoptosis^14,15^, muscle development^16–19^, fertilization^20–22^, unconventional fibroblast growth factor 2 secretion^23^, blood coagulation^24,25^ and a number of immune signaling processes^26–31^. Furthermore, externalization of phosphatidylserine (PS) has been associated with neuronal functions^32^, including synaptic pruning^33,34^, axonal regeneration^35^, and photoreceptor function^36^. Certain mutations in genes encoding scramblases and flippases cause hereditary diseases, including bleeding disorders, hearing loss and progressive familial intrahepatic cholestasis type 1^37–40^. While these findings highlight the importance of lipid asymmetry for eukaryotic physiology, the overarching mechanisms that link trans-bilayer lipid concentration gradients to cell biological function remain unknown: Why is lipid asymmetry required for cell biological function in the first place?

Several hypotheses regarding the cellular functions of lipid asymmetry have been proposed.^3,4,41^ In a biophysical interpretation, the highly ordered outer leaflet could serve as a protective barrier, whereas the more fluid inner leaflet could serve as a platform for catalyzing biochemical reactions. Lipid asymmetry could also regulate lipid-protein interactions by sequestering lipids on opposing membrane leaflets. Finally, lipid asymmetry could constitute a store of potential energy similar to ion or proton gradients across cellular membranes.^2,42^ In this case, lipid flippases and floppases would be equivalent to ion pumps, whereas scramblases would act as ion channel analogs. While all of these hypotheses are intuitively appealing, very little evidence supports any of them.

To comprehensively understand the biology of lipid asymmetry, a systematic approach is required. We designed our study as a multimodal screen for cellular processes affected by genetic perturbation of the lipid asymmetry machinery, followed by a detailed characterization of the most prominent findings. We thus generated a series of ten CRISPR-mediated knockouts (KOs) of lipid flippases, floppases, and scramblases in the plasma membrane of mammalian cells (HCT116). For each KO line, we performed a systems-level analysis of transcriptomic, proteomic, and lipidomic alterations, with large-scale dataset integration to identify shared cellular processes influenced by lipid asymmetry (Fig. 1B). We discovered a broad-spectrum regulation of cellular metabolic pathways extending from transmembrane solute transport to glycolysis, mitochondrial energy transformation, and biosynthesis of lipids, peptides, nucleic acids, and carbohydrates, in addition to previously identified alterations to immune response, cell adhesion, and differentiation. We further validated the role of P_4_-ATPase flippases in energy metabolism by a series of biochemical assays. We found lower steady-state levels and faster consumption of triacylglycerols (TAGs), and higher ATP consumption rates after inhibition of oxidative phosphorylation and glycolysis. Flippase KO lines (ATP11C, ATP8B1, and TMEM30A) showed slower growth, higher mass-specific metabolic rates, and greater heat loss per unit of synthesized biomass, suggesting a shift from anabolic biomass production to a more catabolic state. Altogether, we identify a direct link between lipid asymmetry and cellular metabolism and provide evidence supporting a central role for lipid bilayer asymmetry in cellular energetics.

## Results

### Loss of lipid asymmetry mediators leads to altered lipid trans-bilayer movement

We chose the HCT116 epithelial colon cancer cell line as our model system because it has a near-diploid chromosome number and provides context for the gut-immune system, the gut-brain axis, and metabolism. As targets for genetic modification, we chose plasma membrane-resident lipid translocases and scramblases that we found to be expressed in HCT116 cells: ANO6 and PLSCR1 (scramblases); ABCA1 and ABCA7 (floppases); ATP8A1, ATP11A, ATP11C, ATP8B1, and ATP8B2 (flippases); and the shared flippase subunit TMEM30A (used in our previous study^43^). Most of these proteins are unambiguously confirmed as components of the lipid asymmetry machinery with strong structural and biochemical evidence.^11,39,44,45^

We used CRISPR-mediated deletion of the critical exon present in all protein isoforms for each gene and confirmed knockouts by measuring read coverage for RNA transcripts (Extended Data Fig. 1A). We achieved full deletion of the target exons for all candidates except TMEM30A, which retained a small fraction of the target transcript. We further confirmed this at the protein level, finding that approximately 30% of TMEM30A abundance remained after knockout, whereas other lipid asymmetry proteins were undetectable (Extended Data Fig. 1B). To assess changes in plasma membrane trans-bilayer lipid distribution caused by the genetic perturbations, we used two complementary approaches. We first monitored plasma membrane recruitment of a purified PS biosensor (LactC2-eGFP) as a proxy for increased presence of PS in the outer leaflet using live-cell imaging (Fig. 1C). We found an approximately 30-fold higher LactC2-eGFP signal in TMEM30A KO cells compared with WT cells. TMEM30A is the accessory subunit of many flippase complexes, including the P_4_-ATPase lipid translocases^44,45^, which likely explains the magnitude of the observed effect, which is in line with previous observations.^43,46^ Smaller increases in LactC2-eGFP signal (1.6-to 1.9-fold) were observed in ATP11A, ATP8B2, ATP11C and ABCA7 KO cells, of which only the ATP11A increase was significant. PLSCR1 KO cells showed lower LactC2-eGFP binding than WT cells, whereas the remaining KO lines were very similar to WT (Fig. 1C). Apart from the strong effect for TMEM30A, these data suggest a modest change of PS transbilayer distribution in about half of the KO lines.

To capture changes to lipid transbilayer distribution more generally, we extracted cell surface lipids from WT, TMEM30A and ATP11C KO cells by incubating live cells with methyl-α-cyclodextrin (Fig. 1D), which has a small hydrophobic cavity and therefore preferentially solubilizes phospholipids rather than cholesterol.^47–49^ We analyzed cyclodextrin-complexed lipids as well as the lipidome of the residual cells, which contain all remaining cellular lipids, including those of internal membranes, by shotgun lipidomics to compare relative phospholipid levels in the outer leaflet of the plasma membrane versus the whole cell (Supplementary Table 1). The extracts contained 0.11 to 0.20% of the cellular phospholipids (Extended Data Fig. 2A). The extracts also contained a significant amount of cholesteryl esters (CE), which can largely be explained by residual neutral lipids from fetal bovine serum (FBS) in the culture medium (Extended Data Fig. 2B). We first assessed the levels of PE, PE ether lipids (PE O-) and SM in the lipid extracts, as PE primarily resides in the inner and SM in the outer leaflet of the plasma membrane.^1,2^ Compared with the residual cellular lipids, the extracts of all three cell lines contained lower molar proportions of PE and PE O- and a higher proportion of SM (Fig. 1E), in line with the expectation that methyl-α-cyclodextrin predominantly extracts lipids from the outer leaflet.

As removing outer-leaflet lipids from living cells likely triggers compensatory lipid translocation from the inner leaflet during the 8 min extraction period,^50^ we compared the extracts qualitatively between cell lines rather than interpreting them as the steady-state composition of the outer leaflet. Extracts of both TMEM30A and ATP11C KO cells contained a higher mean molar proportion of PC ether lipids (PC O-) and a lower proportion of PC than extracts of WT cells, and ATP11C KO extracts also contained more SM (Fig. 1E), suggesting that these KOs alter the lipid composition of the outer leaflet. In contrast to the LactC2-eGFP imaging (Fig. 1C), the extracts of TMEM30A KO cells did not contain a higher proportion of PS than those of WT cells (4.6 versus 4.7 mol%). Compensatory translocation during the extraction could reduce such differences between cell lines, so the extracts may underestimate the changes in the outer leaflet. Taken together, these approaches indicate that the genetic perturbations indeed disrupt PM lipid asymmetry.

Next, we assessed the impact of lipid asymmetry knockouts on total cellular lipid levels using quantitative shotgun lipidomics. We monitored major classes of membrane lipids including glycerophospholipids (PS, PE, PC, PI, including ether lipids), sphingolipids (Cer, HexCer and SM), cholesterol and neutral lipids (CE, DG and TAG). We found that PE levels were uniformly elevated, while the molar abundances of TAGs and PC ether lipids were decreased (Fig. 2A-C, Extended Data Fig. 3A). Most other lipid classes were unaffected (Fig. 2A, Extended Data Fig. 3A). Among the main lipid substrates of the lipid translocases in our study, only PE was increased across all KO lines. Among glycerophospholipids, short-chain species (14 to 15 carbon atoms and 0 to 1 double bonds per fatty acid moiety) were enriched regardless of lipid class. This was particularly pronounced for P_4_-ATPase KO lines (Extended Data Fig. 3B). This trend was balanced by changes in longer-chain species, resulting in an overall conservation of the average phospholipid chain length (Fig. 2D, Extended Data Fig. 3B). These compensatory changes were cell line-specific, potentially due to the need to maintain constant bilayer thicknesses. Average length of hydrocarbon moieties and unsaturation were unaffected across membrane lipid classes, whereas changes in neutral lipid content differed between KO lines (Extended Data Fig. 3C). We hypothesized that these species-level changes reflected the substrate specificity of the lipid asymmetry machinery.

**Fig. 2.**
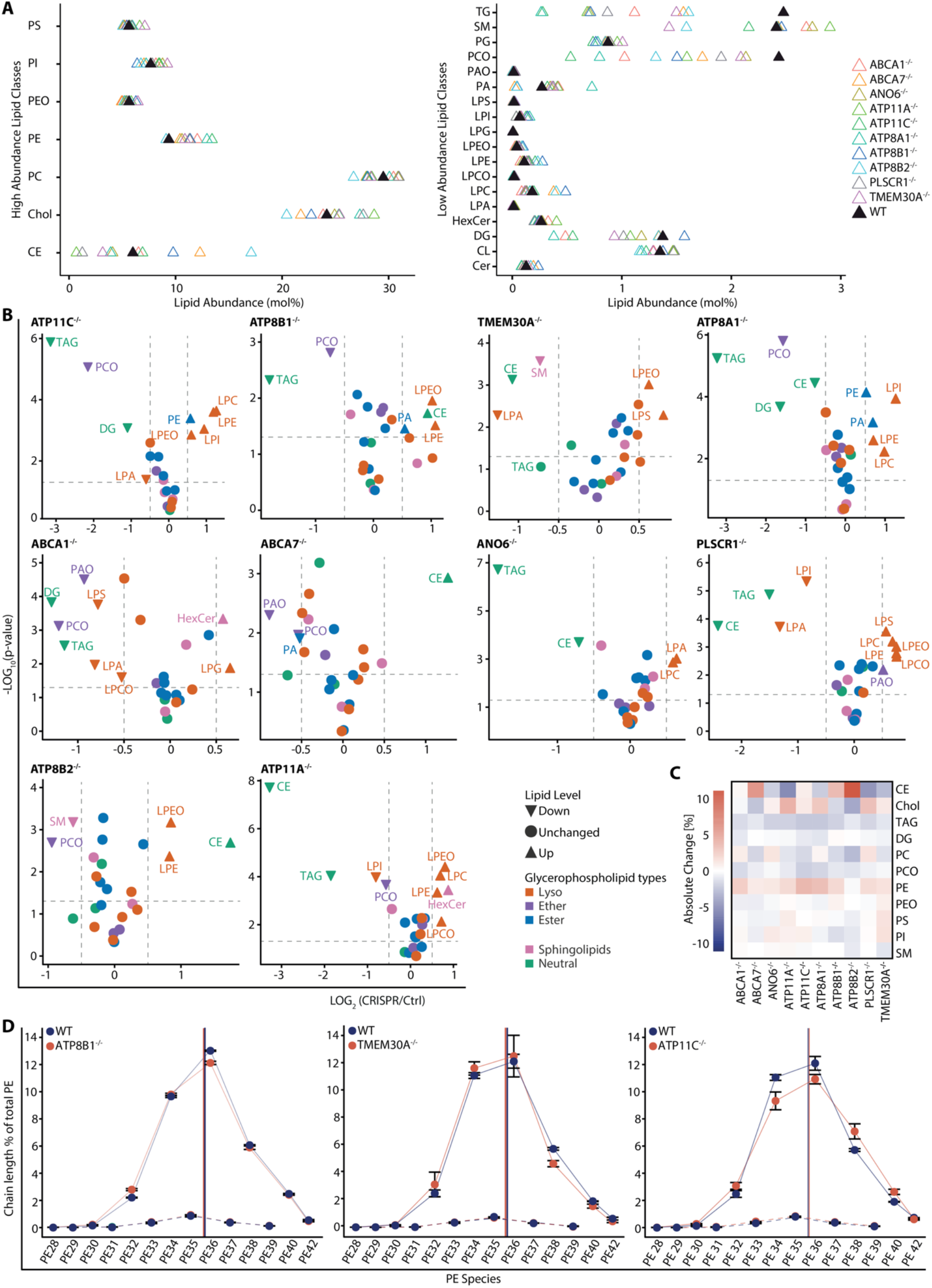
Knockout of lipid translocases alters lipid substrate levels. **(A)** Overall lipid composition of the investigated cells. **(B)** Relative changes of lipid class levels in all KO cell lines compared to WT, shown as volcano plots (x-axis = ratio to control (Log_2_ scale); y-axis = p-value (-Log_10_ scale)). Triangles indicate lipid classes that pass the significance thresholds (dashed lines), pointing up for increased and down for decreased levels, and circlesindicate unchanged classes; colors indicate lipid categories (key at bottom right). Ratios are relative to WT cells of the same measurement set; dashed lines mark a Log_2_ ratio of ±0.5 and p = 0.05. **(C)** Absolute changes in lipid class levels in KO cell lines compared to WT, quantified by lipidomics and shown as a heatmap (key at left). **(D)** Average chain-length of all detected PE species as shown for WT, ATP8B1^-/-^, ATP11C^-/-^ and TMEM30A^-/-^ KOs. Values are the percentage of total PE with a given chain length, multiplied by that chain length and divided by 100, so that they sum to the average chain length (vertical lines). Solid and dashed lines connect species with even and odd chain lengths, respectively. Points show the mean and error bars the SEM of n = 3 (ATP8B1^-/-^ and the corresponding WT) or n = 2 (ATP11C^-/-^, TMEM30A^-/-^ and the corresponding WT) samples. Full list of lipid species in Extended Data Fig. 3B.

To probe changes in lipid dynamics rather than steady-state levels, we measured PE and PC internalization kinetics, as enzymes catalyzing lipid transbilayer movement and intracellular lipid transport are functionally connected (Extended Data Figs. 4-6).^43,51^ Along these lines, discovered that retrograde PE trafficking from the plasma membrane to internal membranes occurs predominantly via non-vesicular lipid transport^43^.

We measured lipid internalization with near-native bifunctional lipid probes functionalized with diazirine and alkyne moieties, which can be photochemically cross-linked to cellular proteins and visualized by fluorescence microscopy (Extended Data Fig. 4A). Bifunctional lipids were incorporated into the outer leaflet of the plasma membrane and internalization monitored in pulse-chase experiments. We compared internalization kinetics for PE(18:1/Y_16_) (with Y denoting a 16-carbon bifunctional fatty acid), which mimics the PE34:1 species that was not altered in our lipidomics data (Extended Data Fig. 3B), and PE(14:0/Y_14_) (with Y_14_ denoting a 14-carbon bifunctional fatty acid), which serves as a model for the medium-chain PE and PS species that are upregulated in the flippase KOs. Using co-stains for the PM and ER, we quantified relative lipid localization over time as the ratio of lipid signal at the PM to the combined signal at the PM and ER at 3, 4, 5, 10 and 30 min after the start of lipid loading (Extended Data Fig. 4, Supplementary Table 2). Interestingly, we observed that plasma membrane labelling with a covalent PM marker dye, was markedly reduced in all three KO lines compared to WT cells at every time point (on average about 2-, 6- and 21-fold lower in TMEM30A, ATP11C and ATP8B1 KO cells, respectively; Extended Data Fig. 4B), which may be an indication of an overall altered organization or composition of the plasma membrane. We also included PC(18:1/Y_16_) as a phospholipid other than PE and PS for comparison. In WT cells, PE(18:1/Y_16_) was transported slightly faster than PE(14:0/Y_14_), and the majority of PE internalization occurred by 10 min (Extended Data Figs. 4D and 5). In ATP8B1, ATP11C, and TMEM30A KOs, both PE species were internalized more slowly than in WT cells and plasma membrane pools remained detectable at all time points tested, with effects of similar magnitude (Extended Data Figs. 4D,E and 5). PC(18:1/Y_16_) was likewise retained at the plasma membrane in the KO lines, although to a lesser extent in TMEM30A KO cells (Extended Data Figs. 4D,E and 6). Taken together, we find that: (i) KO of P4-ATPase protein complex components slowed PE internalization from the plasma membrane of living cells, consistent with reduced flipping, further confirming the functional relevance of the generated cell lines; (ii) this reduction was similar for the long-chain and the medium-chain PE species.

### Multi-omics analysis uncovers a link between lipid asymmetry and metabolism

We performed parallel transcriptomic and proteomic analyses in KO versus WT cells to systematically map cellular processes related to plasma membrane lipid asymmetry. RNA sequencing quantified >25000 genes across all conditions (Supplementary Table 3; see Methods). Whole-cell proteomes were acquired using data-independent acquisition mass spectrometry (DIA-MS), and >7000 proteins were quantified across all KO lines after MS quality filtering (4000-5400 proteins per condition; Supplementary Table 4, Extended Data Figs. 7A-D and 8). By integrating these datasets, we first sought to gain a global view of changes in cellular processes upon perturbation of the lipid asymmetry machinery. We sorted each dataset into six categories via k-means clustering (Euclidean distance), reflecting the average abundance trends (Fig. 3A). Clusters 4 and 5 featured genes that were up-or down-regulated across all CRISPR lines, respectively (for transcriptomics: average 1.7-fold increase or 1.53-fold decrease; for proteomics: average 1.3-fold increase or 1.14-fold decrease). These averages combine all ten KO lines; in single lines, the mean increase of cluster 4 ranged from 1.2-to 2.9-fold for transcripts and from 1.1-to 1.7-fold for proteins (Supplementary Table 5). The mean decrease of cluster 5 reached 2.8-fold for transcripts and 1.3-fold for proteins, and in both clusters the largest changes occurred in P4-ATPase KOs.

**Fig. 3.**
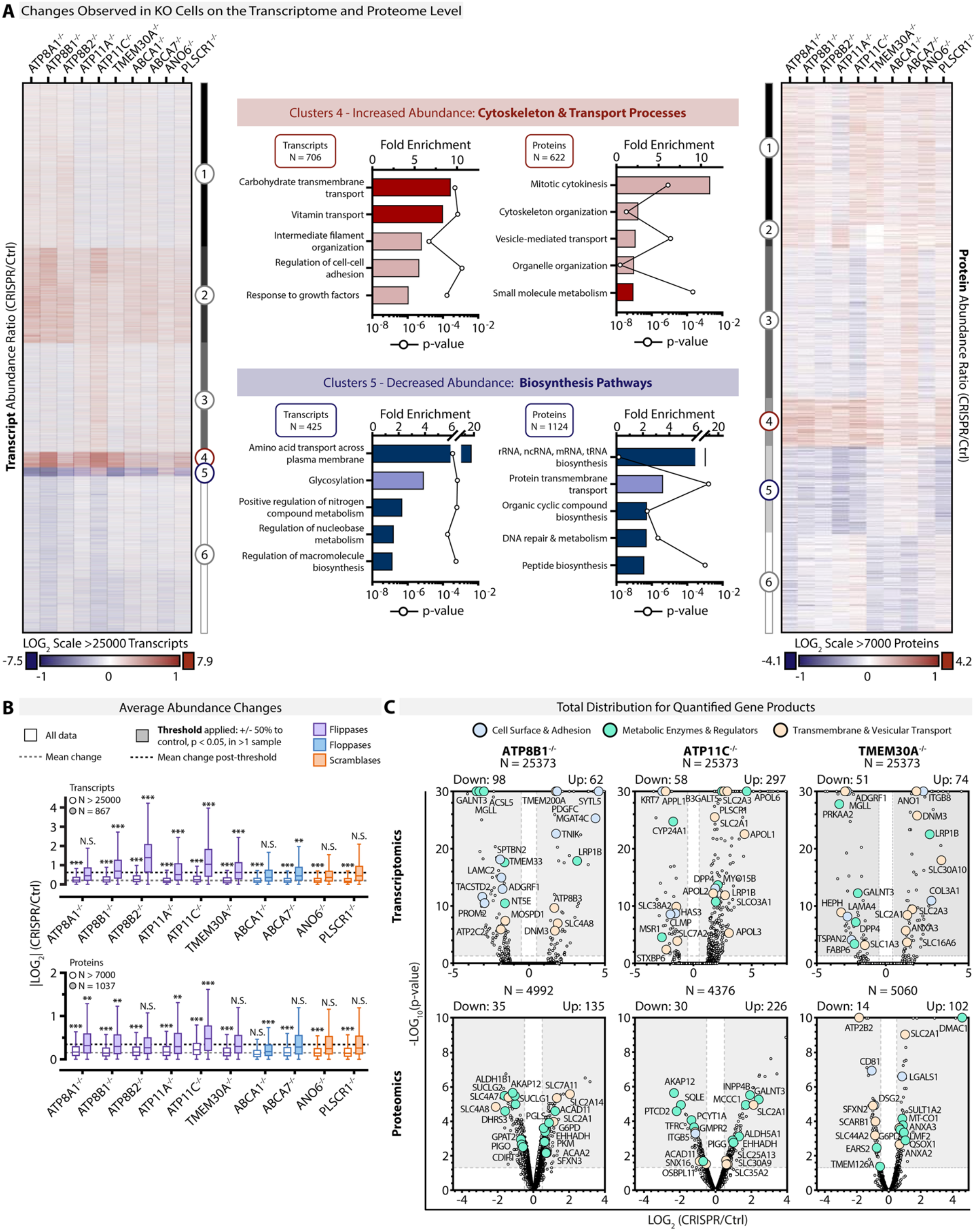
Multi-omics analysis reveals a link between lipid asymmetry and energy metabolism. **(A)** Transcriptomic (left heatmap) and proteomic (right heatmap) dataset heatmaps organized by k-means clustering into differentially regulated genes and proteins across all CRISPR KO lines (indicated at top). Color mapping corresponds to the abundance fold-change to WT (Log_2_ transformed; key below). Gene ontology (middle panel) statistical over-representation test (PantherDB) for Clusters 4 (upregulated across all CRISPR lines) and 5 (downregulated), showing a selected top 5 umbrella terms for each dataset (see Supplementary Table 5 for complete lists). **(B)** Average abundance changes in complete (white boxes) versus post-statistical thresholding (colored boxes) datasets compared to WT, displayed as a box-and-whisker plot (whiskers are Tukey distribution, box line at median; * p < 0.05, ** p < 0.01, *** p < 0.001, N.S. = non-significant for KO versus all data mean as indicated by dashed lines). (C) Volcano plots showing total data distribution for quantified gene products (top = RNA, lower = protein) for ATP8B1, ATP11C, and TMEM30A KOs. Selected points are highlighted by functional category (color key at top) and the gene name is indicated. Shaded regions represent fold-change to WT ≥2 (x-axis) and p < 0.05 (y-axis), with total counts indicated above each graph.

We further analyzed the up- and down-regulated clusters for over-represented gene ontology (GO) terms (Fig. 3A middle panel, Supplementary Table 5). This served two purposes: (i) to test whether the affected genes map to pathways previously linked to lipid asymmetry, providing confidence in our experimental design, and (ii) to identify processes not yet linked to lipid asymmetry. We chose to retain moderately regulated genes in the pathway-level analysis to boost bioinformatic identification of biological processes, because pathway regulation can involve large changes in a few enzymes and moderate or no changes in others, or shifts in the molar ratios between competing enzymes.^52^ Individual genes were then assessed separately with stricter thresholds (±50% change to control, p < 0.05; Fig. 3B,C) to select candidates for follow-up experiments.

Because lipid asymmetry is a core feature of membrane organization, disrupting it is expected to affect several processes. Many GO terms enriched in the transcript and proteome datasets included cellular processes previously linked to plasma membrane lipid asymmetry, expanding evidence for multiple context-specific studies and supporting our experimental design. Genes with increased abundance (Clusters 4) are involved in cytoskeleton organization, cell adhesion, and membrane trafficking (Fig. 3A), in agreement with the known functions of lipid asymmetry in cell adhesion and migration in mammalian cells^53–57^ and membrane trafficking in yeast^58–62^. On the other hand, downregulated genes (Clusters 5) are enriched for GO terms related to cell surface signaling and neurological processes (Supplementary Table 5), reflecting the prevalence of flippase mutations in neurological disorders (e.g., ATP8B1).^37–39^ In the transcriptomic data, Cluster 4 also included upregulated immune response GO terms, consistent with a reported role for ATP11C in B-cell maturation and macrophage function.^63–65^

The upregulation of genes involved in membrane trafficking led us to test for alterations in membrane trafficking. P_4_-ATPases have been implicated in endocytosis in yeast, and here we found that transcripts of endocytosis-associated genes were overall reduced in the ATP8A1, ATP8B1 and ATP8B2 KOs and increased in the TMEM30A KO, whereas at the protein level only the ATP8B1 KO showed a corresponding shift (Extended Data Fig. 9A). We therefore measured the uptake of the protein cargo transferrin, as well as the number and size of transferrin-positive structures in the ATP11C, ATP8B1 and TMEM30A KO lines used for the lipid transport experiments (Extended Data Fig. 9B,C). All three KO lines internalized transferrin, with modest changes in uptake and in the number, size and intensity of transferrin-positive structures. While the mean intensity of individual transferrin puncta after 30 min was lower in all three KO lines, other readouts changed in a gene-specific manner, in some cases in opposite directions. For example, the size of the endocytic recycling compartment was smaller in the ATP11C KO and larger in the ATP8B1 and TMEM30A KOs, and total transferrin per cell was lower in the ATP11C KO only (Extended Data Fig. 9C). In contrast to non-vesicular lipid transport of PE, the effects on vesicular trafficking are thus largely gene-specific, suggesting that they reflect distinct roles of the respective P_4_-ATPases in vesicular transport.

A more striking trend was that many of the over-represented GO terms represented numerous aspects of metabolism. This was apparent at both the RNA and protein levels and in both upregulated and downregulated genes, indicating a multifaceted impact of lipid asymmetry perturbations on cellular metabolism. Specifically, carbohydrate and vitamin transmembrane transport were among the top enriched GO terms for upregulated transcripts, complemented by small molecule metabolism in the proteomics dataset (Clusters 4; Fig. 3A). In gene products with decreased abundance, we found enrichment of numerous GO terms in both datasets for general metabolic pathways ranging from amino acid transport to molecular biosynthesis (e.g., nitrogen compound, nucleobase, heterocycle, amide), macromolecular biosynthesis (DNA, m/r/t/nc/snRNA, peptide), and cellular component biogenesis (Clusters 5; Fig. 3A; Supplementary Table 5). Overall, loss of lipid asymmetry appears to shift soluble nutrient transport from amino acids to carbohydrates and induce a corresponding downregulation of macromolecular biosynthesis.

While this general trend is present in all KO lines, we noted that the individual hits passing our significance thresholds differ. Therefore, we analyzed gene-specific trends to narrow down targets and processes for functional follow-ups. First, we found that the most robust changes were caused by KO of P_4_-ATPase flippases, in both the pre- and post-thresholded data (Fig. 3B), which also showed high correlation for transcript and protein abundance changes among all family members (Extended Data Fig. 8B). In contrast, the responses of the P4-ATPase flippase KOs and of the scramblase KOs overlapped only partly (Extended Data Fig. 10). The gene-wise median changes of the five P_4_-ATPase KOs correlated only weakly with those of the ANO6 and PLSCR1 scramblase KOs on the transcript and protein level (Spearman ρ = 0.28 to 0.35). Fewer than 3% of transcripts and 0.2% of proteins changed at least 1.5-fold in both, mostly in the same direction (Extended Data Fig. 10A,B). The two scramblase KOs correlated even more weakly with each other (Spearman ρ = 0.12 for transcripts and 0.19 for proteins), and 443 transcripts (1.7%) and 6 proteins (0.3%) changed at least 1.5-fold in both (Supplementary Tables 3 and 4). Of the 1651 transcripts whose median change in the P_4_-ATPase KOs reached the 1.5-fold threshold, 736 changed in neither scramblase KO, and only 157 changed in the same direction in both (Extended Data Fig. 10C). At the protein level, 7 of the 11 proteins that reached this threshold in the P4-ATPase KOs changed in neither scramblase KO (Extended Data Fig. 10D). Second, among the genes passing our significance cut-offs in the P_4_-ATPase KOs, we found many representative members of the pathways identified by GO enrichment, especially transmembrane solute carriers (SLCs) that exhibited some of the most robust changes in every dataset (Fig. 3C). The changes of many of these hits were considerably larger than the average changes of their clusters. The broad metabolic regulation was most apparent at the protein level, with common hits including the glucose transporter GLUT1 (SLC2A1; average 3.5-fold increase in ATP11C, ATP8B1 and TMEM30A KOs), the pentose phosphate pathway enzymes glucose-6-phosphate dehydrogenase (G6PD; average 1.7-fold increase) and 6-phosphofructokinase (PFKL; average 1.5-fold increase in ATP8B1 and TMEM30A KOs), and the pyruvate kinase (PKM, average 1.6-fold increase in ATP11C and ATP8B1 KOs) that catalyzes the final rate-limiting step of glycolysis. Similarly, proteins involved in peroxisomal and mitochondrial beta-oxidation, such as the peroxisomal bifunctional enzyme (EHHADH) and the mitochondrial carnitine O-palmitoyltransferase 2 (CPT2), are increased in ATP11C and ATP8B1 KOs (Fig. 3C). Furthermore, the metabolic shift from amino acid to carbohydrate membrane transport is apparent in both data sets, as most metabolic GO terms were in the downregulated clusters. Based on these findings, we decided to carry out a mechanistic characterization of the link between maintenance of lipid asymmetry and regulation of cellular metabolism, focusing on the P_4_-ATPases.

### Perturbing lipid asymmetry changes global lipid composition and increases neutral lipid consumption

Among the metabolic hits, we noted that gene products mapping to lipid biosynthesis, beta-oxidation, and lipid droplet maintenance pathways changed in abundance at both the protein and mRNA levels (Figs. 3 and 4A,B). Examples passing our significance cut-offs in multiple KOs include upregulation of the ER-anchored GPI ethanolamine transferase 2 (PIGG; 2-fold average increase), the peroxisomal enoyl-CoA dehydrogenase (EHHADH; 1.8-fold), and the mitochondrial long-chain fatty acid transporter CPT2 (1.5-fold), which were all shared by ATP11C and ATP8B1 KOs. Downregulation of the lipid droplet short-chain dehydrogenase/reductase 3 (DHRS3; average 2.1-fold decrease) was shared between ATP8B1 and TMEM30A KOs, while mitochondrial glycerol-3-phosphate acyltransferase 2 (GPAT2; 1.8-fold decrease) and the ER-anchored very long chain fatty acid elongase 7 (ELOVL7; 1.4-fold decrease) passed significance thresholds in ATP8B1 KO, among others (see Supplementary Table 6 for complete GO-annotated hits in all KO lines). Again, we turned to quantitative shotgun lipidomics to test whether these changes are reflected in cellular lipidome homeostasis. Among various KO-specific trends, we found that PE, lysophosphatidylethanolamine (LPE) and ether lysophosphatidylethanolamine (LPEO) were elevated in nearly all KOs (Fig. 2B,C and Extended Data Fig. 3A; Supplementary Table 7). In contrast, levels of triacylglycerol (TAG) and ether phosphatidylcholine (PCO) decreased. TMEM30A also featured downregulated levels of the second major neutral lipid class, cholesterol esters (Fig. 2B). Increased PE is likely an adaptation to perturbed lipid translocase levels, as PE and PS are substrates of P_4_-ATPases, which is in-line with our observation that PE internalization was slower upon KO (Extended Data Figs. 4D,E and 5). On the other hand, TAGs (decreased) are primary energy storage lipids and major components of lipid droplets, and lysophospholipids (increased) are the product of fatty acid mobilization from glycerophospholipids. Together, these observations suggest increased fatty acid flux toward beta-oxidation for ATP generation. To test for altered TAG consumption rates, we conducted an oleic acid feeding experiment that induces enlarged lipid droplets. We then monitored lipid droplet shrinkage by fluorescence microscopy, which indicates neutral lipid consumption. We found that initial lipid droplet formation was significantly reduced in ATP11C KO cells compared to WT, whereas ATP8B1 and TMEM30A KOs exhibited similar trends (Fig. 4C,D). However, all three KO lines exhibited faster shrinking of lipid droplets after removal of oleic acid (Fig. 4C,D), indicating higher neutral lipid consumption, while maintaining a similar number of LDs (Extended Data Fig. 7E). Taken together, our findings suggest that maintenance of lipid asymmetry involves cellular lipid storage pathways.

**Fig. 4.**
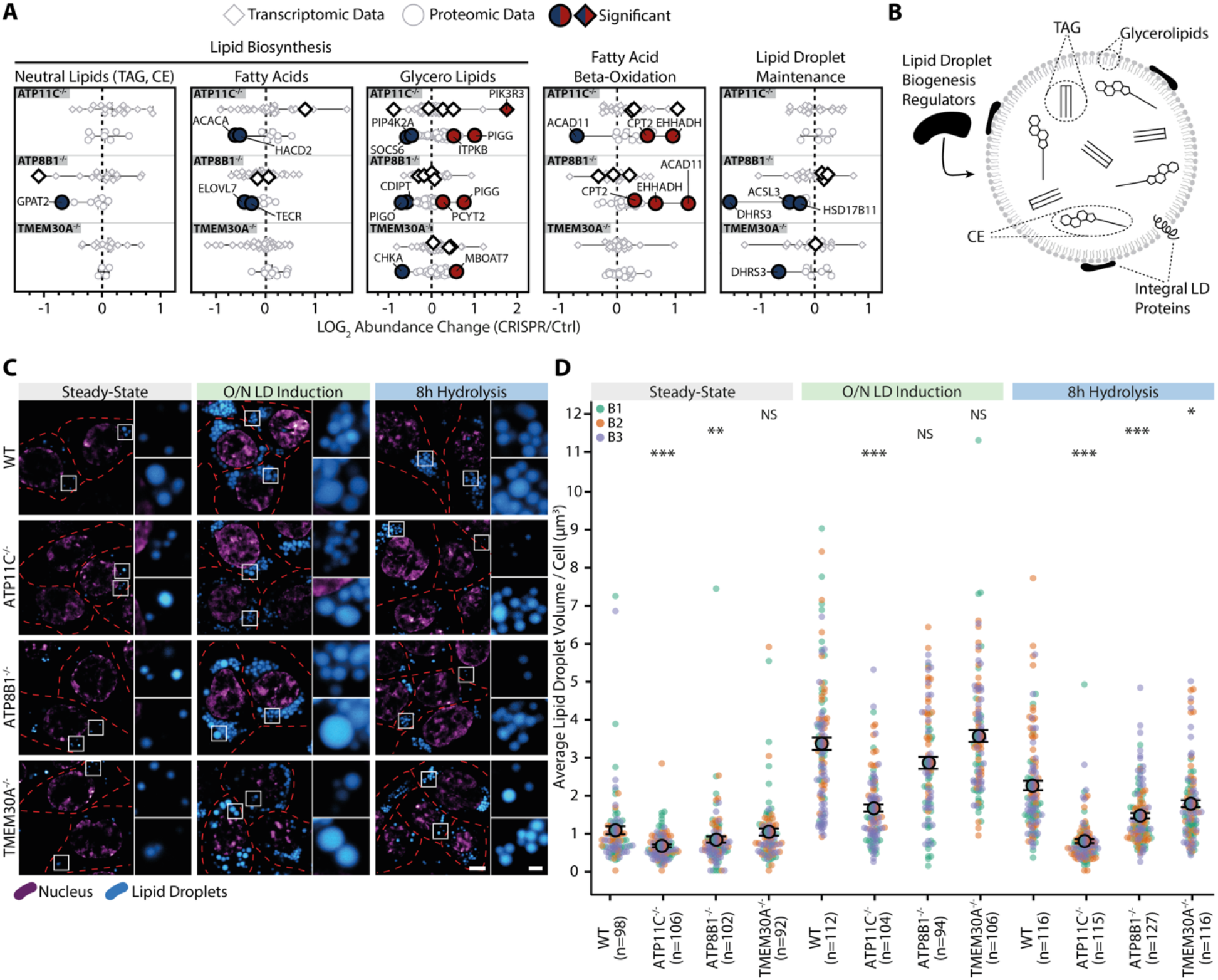
Loss of flippase function leads to faster neutral lipid hydrolysis rates. **(A)** Proteins and transcripts related to lipid metabolism are differentially regulated in P_4_-ATPase KOs. Plotted are the abundance ratios to control cells (Log_2_ scale) for all genes corresponding to that group, sorted by KO line (top to bottom). Each point is an individual gene/protein, with selected genes that pass p<0.05 in at least one omics dataset per KO labeled, enlarged, and colored in the dataset passing significance. See also Supplementary Table 6. **(B)** Schematic overview of lipid droplet (LD) composition. **(C)** Assessment of LD hydrolysis in WT versus KO cell lines following overnight (O/N) incubation with oleic acid to induce LD generation (O/N LD induction) and after a further 8 h in fresh medium (8h hydrolysis). LDs were visualized using LipidSpot, and Hoechst for the nucleus. Red dashed lines indicate cell boundaries, one mid-plane z-slice is shown. Scale bars = 5 µm and 1 µm in the insets. **(D)** Average LD volume per cell, quantified from images as in panel C. All points are shown, colored by biological replicate (B1-B3), with n = number of analyzed cells, p-values above (two-sided Mann-Whitney-U test for each KO compared to WT), and error bars indicating standard error of the mean. p-Values: *** < 0.0001, ** < 0.001, * < 0.01, NS = not significant.

### Knockout of P_4_-ATPases alters cellular bioenergetics

Our proteomic, transcriptomic, and lipidomic datasets point to a global shift in energy metabolism driven by altered lipid trans-bilayer dynamics. Significantly altered gene abundances were observed for major metabolic pathways across subcellular space, including input (transmembrane transport), cytosolic processing (glycolysis, pentose phosphate), and mitochondrial energy output (TCA cycle) (Fig. 5A). Specifically, we found numerous hits related to cellular nutrient uptake by active transmembrane transport, such as GLUT1 (SLC2A1) which we further confirmed by immunofluorescence (Fig. 5B, Extended Data Fig. 11A). The enzymes catalyzing the rate-limiting steps of both the pentose phosphate (glucose-6 phosphate1-dehydrogenase, G6PD; average 1.75-fold increase) and glycolysis (pyruvate kinase, PKM; average 1.6-fold increase) pathways are among the top upregulated proteins across all P_4_-ATPase KOs (Fig 5A). Additionally, we observed significant changes in TCA cycle proteins (Fig. 5A) and beta-oxidation machinery (Fig. 4A), along with decreased TAG levels and elevated neutral lipid consumption (Figs. 2B,C and 4C,D).

**Fig. 5.**
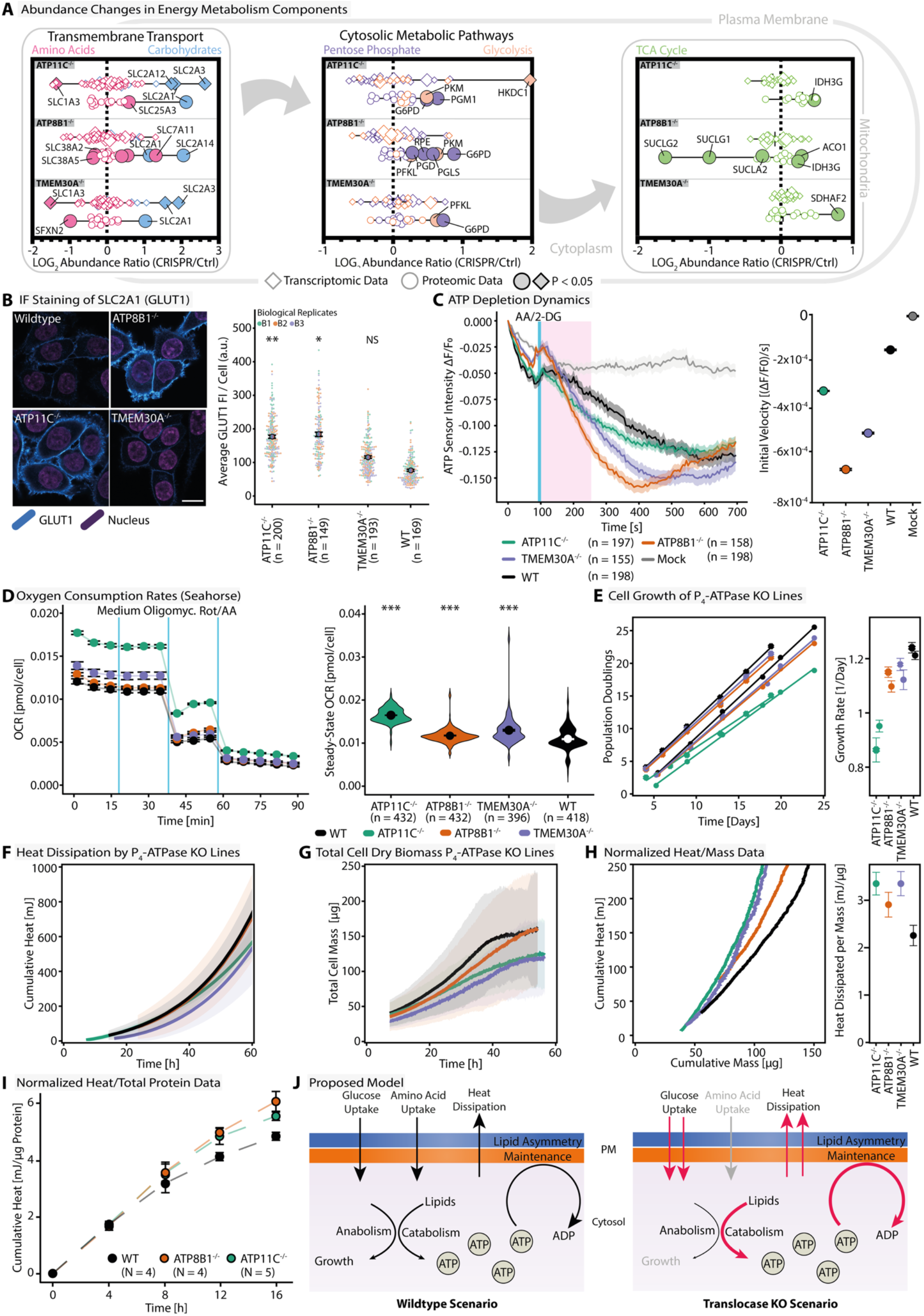
Loss of flippase function leads to inefficient production of biomass. **(A)** Quantified genes corresponding to major cellular metabolic processes, organized by subcellular location. Each plot is the abundance ratio to control (Log_2_ scale) for all genes in the datasets corresponding to that process, sorted by KO line (top to bottom) and colored by pathway. Selected genes that pass p<0.05 in at least one omics dataset per KO are labeled, enlarged, and dark-colored for the dataset passing significance. See also Supplementary Table 6. **(B)** Immunofluorescence staining of endogenous GLUT1 (SLC2A1) in WT versus KO cells (left panel) and analysis of mean fluorescence intensity (FI) per cell (right panel). Scale bar is 15 µm. Coloured points show single cell measurements (n values given on x-axis) with colours indicating biological replicates, black circles show averages and error bars indicate standard error of the mean. Statistical analysis was performed using Student’s t-test for each KO compared to WT. GLUT1 staining of other KO lines can be found in Extended Data Fig. 11A. **(C)** Depletion of cellular ATP measured with a fluorescent ATP sensor (iATPSnFR) after the addition of Antimycin A and 2-Deoxy-D-glucose (2-DG), at the timepoint indicated by the blue line (average traces are shown, n = number of analyzed cells, shading is standard error of the mean). The area shaded in pink indicates the timeframe used for the calculation of initial velocities (right panel; error bars are standard error of the mean). **(D)** Assessment of cellular oxygen consumption rates (OCRs) via Seahorse assay. Left panel: OCR measurements before and after the addition of medium, Oligomycin and Rotenone (Rot)/Antimycin A (AA), indicated by the blue lines. Right panel: steady-state OCR of the investigated cell lines shown as a violin plot (n = number of total measurements from 22 to 24 wells per cell line in each of three independent experiments, pooled over measurement cycles and normalized to the number of cells per well; p-values calculated via two-sided Mann-Whitney-U test of KO versus WT; circle is mean; error bars are standard error of the mean). **(E)** Cell growth expressed in cumulative population doublings, inferred from cell counting during routine passaging (left panel), and corresponding growth rates calculated from linear regression (right panel). Symbols denote two independent batches. Error bars are standard deviation of growth rates bootstrapped distribution with 10000 resamplings. **(F)** Cumulative heat dissipated by KO lines. Shading is the standard deviation of 3 biological replicates. **(G)** Total cell dry biomass measured by quantitative phase microscopy. Shading is standard deviation of biological replicates (WT = 5, ATP11C^-/-^ = 4, ATP8B1^-/-^ = 5, TMEM30A^-/-^ = 3). **(H)** Dissipated cumulative heat normalized by cumulative cell dry biomass (time-matched data from Fig. 5F and 5G, left panel). Slopes of linear regression fits of mass-normalized heat data, quantifying the heat dissipation per µg of cell dry biomass grown (right panel). Error bars of right panel are standard deviation of fit slopes bootstrapped distribution with 10000 resamplings including random 20% of original datapoints. **(I)** Cumulative heat dissipated by P_4_-ATPase cell lines normalized to total protein as determined by calorimetry (N = number of biological replicates; error bars are standard error of the mean; dashed lines are linear interpolations). **(J)** Schematic representation of altered bioenergetics after disturbance of plasma membrane lipid asymmetry in WT and KO scenarios. p-Values: *** < 0.0001, ** < 0.001, * < 0.01, NS = not significant.

We therefore predicted that disrupting lipid asymmetry at the plasma membrane would alter total cellular bioenergetics. We used several orthogonal methodologies to characterize the putative changes in global energy metabolism. First, measurements of steady-state levels of ATP, ADP, and the ATP/ADP ratio showed minor differences between cell lines but no clear trends (Extended Data Fig. 11B). We thus assayed the dynamics of cellular energy metabolism with time-resolved assays. Using a fluorescent ATP biosensor, we found faster ATP depletion kinetics for ATP11C, ATP8B1, and TMEM30A KOs compared to WT after inhibition of both oxidative phosphorylation and glycolysis (Fig. 5C). Furthermore, all tested KO lines exhibited faster oxygen consumption rates (OCR), specifically 50 ± 2% for ATP11C, 7 ± 1% for ATP8B1, and 18 ± 2% for TMEM30A (Fig. 5D, Extended Data Fig. 11C,D).

Increased metabolic activity could be related to faster cell growth. However, cell counting showed that flippase KOs lines grow slower than WT cells (Fig. 5E). We hypothesized that disruption of lipid asymmetry may lead to an imbalance between cellular energy expenditure and growth, shifting nutrient resources from a more anabolic growth-promoting to a more catabolic energy-consuming metabolic state. Thus, we may expect that the observed metabolic changes in KOs lines lead to elevated energy expenditure per unit biomass grown. To test this, we measured cellular energy expenditure in the form of dissipated heat using isothermal microcalorimetry (Fig. 5F). In parallel, we quantified cellular growth by measurements of total cellular dry mass accumulation using quantitative phase microscopy (Fig. 5G). Quantitative phase microscopy measures the phase shift of light passing through the cells, which is proportional to their dry mass, and thus quantifies biomass without labeling.^66,67^ KO lines showed reduced heat dissipation and biomass accumulation compared with WT cells, consistent with our cell-counting results. To investigate whether KO lines expend more energy for growth, we plotted the dissipated heat against the accumulated total cell mass (Fig. 5H, left panel). Using data from the exponential growth phase, we found that heat and mass are linearly correlated for all cell lines measured. The slope quantifies the cellular energy expenditure required to synthesize 1 µg of cellular biomass. Linear regression showed that all KO lines dissipated more heat per biomass than WT cells (Fig. 5H right panel). Similarly, normalizing heat dissipation by total protein content showed that KO lines dissipated more heat per protein made (Fig. 5I, Extended Data Fig. 11E). These data show that disrupting lipid asymmetry leads to slower but more energy-intensive cellular growth.

## Discussion

Here, we report a systems-level investigation into the broader biological role of lipid asymmetry in cells. Lipid asymmetry has been previously linked to various specific cellular processes and biophysical properties of cellular membranes. Indeed, we found a number of affected processes previously linked to lipid asymmetry, ranging from immunity^26–31,63–65^, cytoskeleton organization^54^, to neurobiology^32–36^ (Fig. 3). In addition, we discovered that lipid asymmetry and cellular metabolism are connected by assessing transcriptome, proteome, and lipidome changes caused by genomic perturbation of the protein machinery associated with lipid asymmetry (flippases, floppases and scramblases). Although cellular lipidome changes were relatively minor, transcriptomic and proteomic analyses revealed changes in global metabolic GO terms, indicating broader rewiring of cellular metabolism. These over-represented metabolic GO terms were particularly apparent in the P_4_-ATPase flippase KO lines. They included down-regulated genes involved in amino acid uptake, peptide and macromolecular synthesis, and up-regulated genes facilitating carbohydrate and vitamin uptake (Fig. 3). Specifically, upregulation of genes involved in beta-oxidation, carbohydrate uptake, glycolysis and the pentose phosphate pathway (Figs. 4 and 5) was accompanied by faster neutral lipid consumption, increased respiration, and ATP usage despite slower cell growth and heat dissipation. Normalizing these multifaceted metabolic changes to cell growth revealed that cells with perturbed lipid asymmetry are characterized by increased heat dissipation or energy expenditure per unit biomass or protein synthesized. Thus, disruption of the lipid asymmetry machinery leads to global metabolic and energetic effects that manifest in slower but more energy-intensive cellular growth.

Our data are consistent with a model in which perturbations of plasma membrane lipid asymmetry cause a global rewiring of cellular nutrient uptake, processing, and energy transduction. KOs of flippases shift nutrient uptake from balanced sugar and amino acid uptake that supplies energy and precursors for anabolic growth to reduced amino acid uptake and a more dissipative, energy-consuming, catabolic metabolism fueled by sugars and fats, characterized by slower cell growth (Fig. 5J). Specifically, KO cells dissipate more energy to produce biomass despite slower growth, highlighting a shift from an anabolic to a more catabolic metabolism. Two explanations for this shift are consistent with our data, and they are not mutually exclusive.

First, KO cells might show more futile cycles as other translocases move nonoptimal substrates across a concentration gradient to maintain lipid asymmetry, increasing ATP consumption and energy dissipation per successful lipid translocation event. Assuming that cells in culture grow at their maximum metabolic capacity, this increased ATP demand to ensure functional asymmetric membranes and viability may come at the cost of reducing other energy-demanding cellular processes. KO cells show down-regulation of genes associated with macromolecule and peptide synthesis and reduced amino acid uptake. This suggests that slower growth results from reduced ribosomal biogenesis and protein translation, which can account for up to 60% of the cellular energy budget in growing cells.^68^ Thus, the observed global metabolic rewiring may be a compensatory response that budgets cellular energy expenditure to meet the elevated energy demands of maintaining functional asymmetric membranes by slowing energy-expensive macromolecular synthesis and growth. Our finding that the loss of individual flippases can affect overall cellular energetics initially seems surprising, as it suggests that maintaining baseline lipid asymmetry places significant demands on the cellular energy budget. This is only possible if high rates of active lipid trans-bilayer movement persist at steady state, along with the corresponding ATP hydrolysis rate.

Second, it is possible that lipid asymmetry also fulfills a similar role to other transmembrane concentration gradients (ions, protons) as a store of potential energy built by pump-like proteins (flippases) to provide energy required for coupled transmembrane processes. Like uncoupling proteins, channel-like proteins (scramblases) can dissipate the concentration gradients^69^, potentially regulating the magnitude of stored energy in asymmetric membranes. This interpretation is supported by membrane reshaping/remodeling observed in response to scrambling during molecular dynamics simulations^69,70^ and in-vitro model membrane experiments^71^. We cannot rule out that some solute carrier proteins facilitate nutrient uptake by secondary active transport driven by a gradient of asymmetrically distributed lipids, and that the substantial remodeling of the SLC proteome observed in KO cells is an adaptation to a perturbed lipid environment. Furthermore, it has been proposed that lipid translocases drive directional transport of lipids between organelles in combination with bridge-like lipid transfer proteins^72^, a model for which we have recently provided further experimental evidence^43^. This in turn suggests that the asymmetric lipid distribution of late secretory pathway organelles could be the energy reservoir required for directional non-vesicular lipid transport between organelle membranes, which is increasingly seen as the primary process for maintaining functional organelle lipid composition^43^. The slower lipid transport from the plasma membrane to the ER in the P_4_-ATPase KOs is consistent with this model (Extended Data Figs. 4-6), although this transport does not by itself report on ATP consumption. Our data do not distinguish between a role of lipid asymmetry as an energy store and the energetic cost of compensatory lipid translocation, and both may contribute. Taken together, our findings pave the way to linking lipid asymmetry, lipid transport, organelle membrane identity, and cellular bioenergetics.

Our approach has two limitations. Direct measurements of lipid asymmetry in nucleated cells remain challenging, and our proxies report changes in PS exposure and in extractable outer leaflet lipids rather than the steady-state composition of both leaflets. In addition, the link between lipid asymmetry and cell energetics rests on genetic perturbation of the translocase machinery, so functions of individual translocases beyond asymmetry maintenance may also contribute to the energetic phenotypes. How the altered lipid environment affects the activity of nutrient transporters such as GLUT1, and how much ATP lipid translocation consumes, are important questions for future work.

## Author Contributions

PB and AN conceptualized and designed the project. AS, JR and AN supervised the project. PB prepared figures with contributions from KCC, VB and SMK. PB prepared samples for all omics analyses with contribution from SMK. TJ performed proteomics experiments. KCC, PB, TJ and AN analyzed transcriptomic and proteomic data. KB synthesized lipid probes. KCC, PB and HML performed lipid imaging experiments. PB and TRR performed cyclodextrin extractions. KS and SR performed lipidomics experiments. KS and PB analyzed lipidomics data. PB analyzed cyclodextrin and lipid imaging data. PB, SMK and CJL performed fluorescence microscopy experiments. PB analyzed fluorescence microscopy data with contribution of SMK. PB performed Seahorse assays. PB and TCA analyzed Seahorse data. ST performed ATP/ADP measurements. ST and PB analyzed ATP/ADP data. VB performed the quantitative phase microscopy experiments and analyzed data. PB, VB and JR performed calorimetry experiments and analyzed data. PB and AN wrote the manuscript with contribution of KCC, VB and JR. All authors read and commented on the manuscript.

## Acknowledgements

This work was supported by the Max Planck Society. AN and JR gratefully acknowledge financial support by the European Research Council (ERC) under the European Union’s Horizon 2020 research and innovation program (grant agreements no GA 758334 ASYMMEM and GA 949811 EnBioSys). AN and AS acknowledge financial support by the Deutsche Forschungsgemeinschaft (DFG) via the TRR83 consortium. This research was supported by an Allen Distinguished Investigator Award, a Paul G. Allen Frontiers Group advised grant of the Paul G. Allen Family Foundation to AN. We thank the following services and facilities at MPI-CBG Dresden for their support: Protein Expression Facility, Scientific Computing Facility, Genome Engineering Facility, Technology Development Studio and the Light Microscopy Facility. We thank Christina Eugster and Eva Angermann from the MPI-CBG Organoid and Stem Cell Facility for their support with cell culture. Furthermore, we would like to thank the Genome Center at CRTD/CMCB for their support.

## Code and data availability

All data and code used in this study can be accessed on the following repository: EDMOND (https://doi.org/10.17617/3.1HISIJ).

## Extended Data Figures

**Extended Data Fig. 1.**
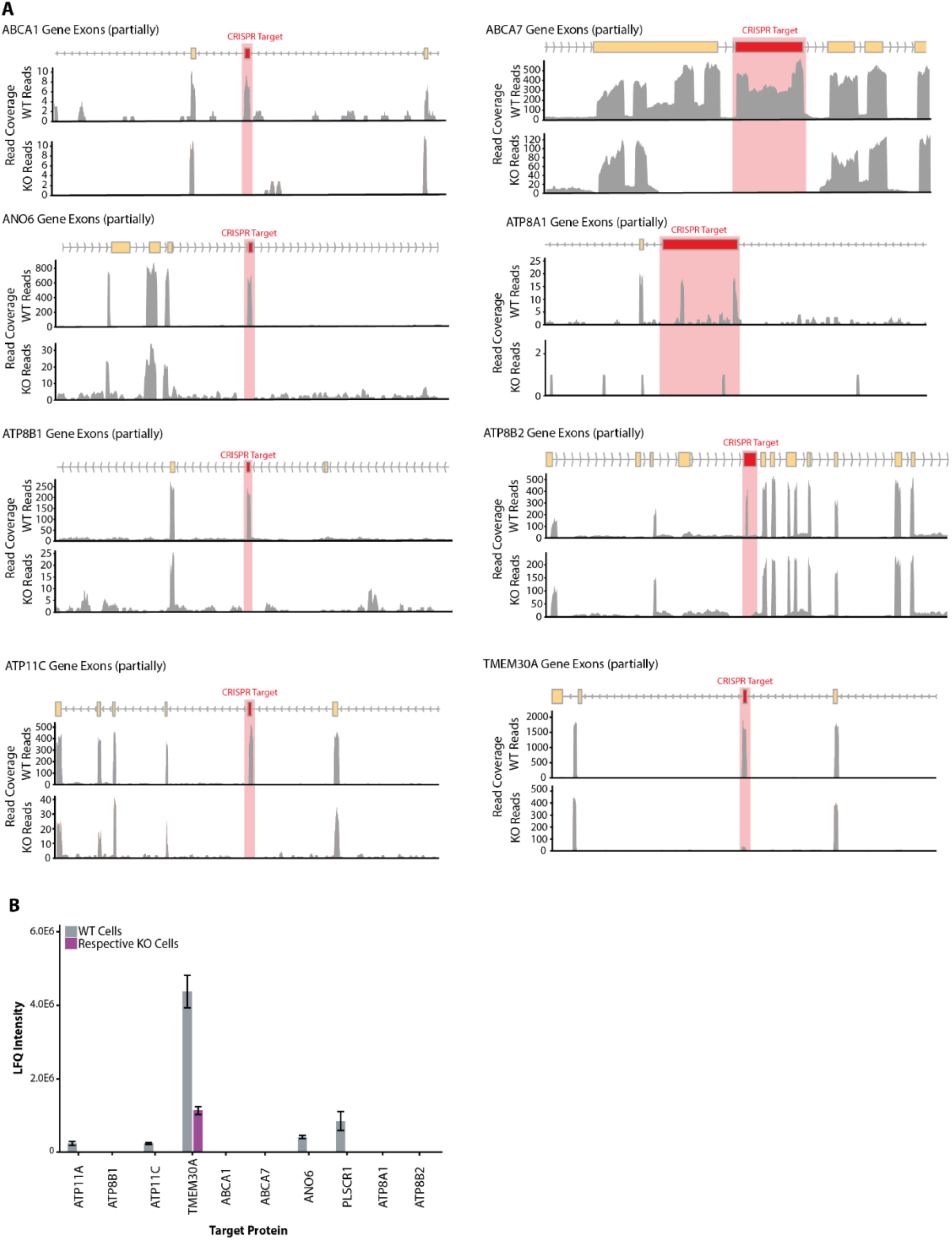
Confirmation of the successful deletion of the target exon. **(A)** Read coverage of the partial gene regions for all KO targets that were close to the targeted exon. WT and KO reads of the respective regions are shown and the red bar indicates the targeted exon for deletion. **(B)** Proteomic assessment of targeted proteins for KO. For ATP11A, ATP11C, TMEM30A, ANO6 and PLSCR1 protein was detected in the WT cells (grey bar) and the amount of protein detected in the respective KO shown in violet. Error bars indicate standard error of the mean.

**Extended Data Fig. 2.**
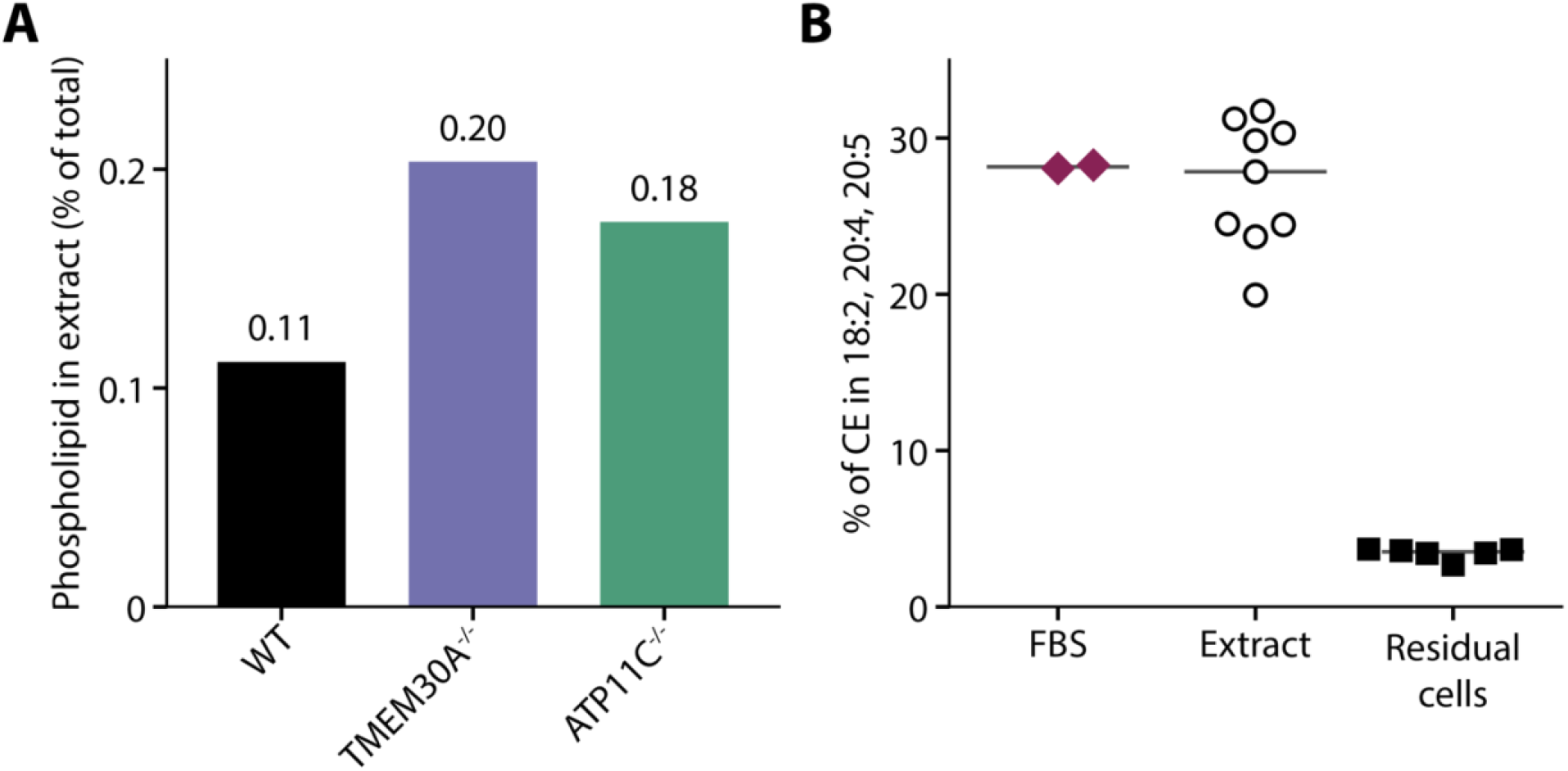
Phospholipid yield and serum carry-over following cyclodextrin extraction. HCT116 WT and KO cells were incubated with 4 mM unloaded methyl-α-cyclodextrin for 8 min at room temperature, followed by lipidomic analysis of the collected supernatants and residual cells. **(A)** Phospholipid in the extract as a percentage of the phospholipid in extract and residual cells together, for the nine measured classes (PC, PC O-, PE, PE O-, PS, SM, PG, LPC and LPE), calculated from separate means for each fraction, using nominal volumes of 3 mL for extracts and 2 mL for residual cells to scale the amounts measured in 200 µL aliquots. **(B)** Share of cholesteryl ester (CE) in the species CE 18:2, CE 20:4 and CE 20:5, which are enriched in fetal bovine serum (FBS) relative to the cells (see Methods), in FBS, extracts and residual cells. Points represent samples (diamonds, the 2 µL and 20 µL FBS samples; open circles, extracts; filled squares, residual cells) and horizontal lines indicate medians. Species with an odd number of carbon atoms were excluded, and phospholipid species of the extracts were kept only if they were also detected in the residual cells. PS in the residual cells is likely underestimated (see Methods). Technical injections were averaged before analysis, with n = 3, 3 and 3 extract samples and n = 1, 2 and 3 residual-cell samples for WT, TMEM30A and ATP11C, respectively; FBS was measured once at 2 µL and twice at 20 µL. Extracts and residual cells were filtered separately.

**Extended Data Fig. 3.**
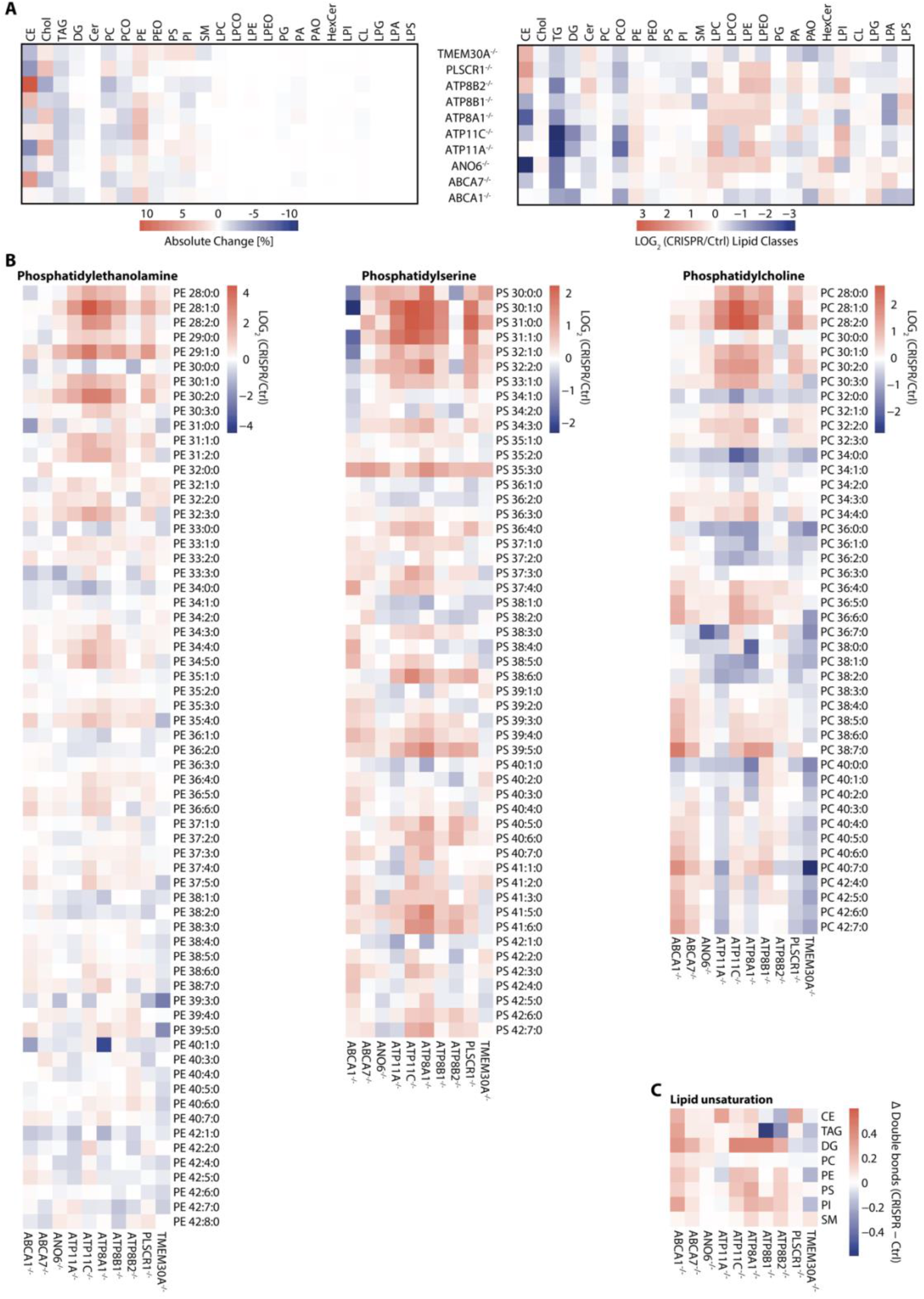
Analysis of shotgun lipidomics data. **(A)** Absolute (left panel) and relative (right panel, Log_2_ scale) lipid class changes in all KO cell lines compared to WT. **(B)** Relative changes in lipid species levels of PE, PS and PC in KO cells compared to WT. **(C)** Changes in the mean number of double bonds per lipid species within each lipid class in all KO cell lines compared to WT of the same measurement set, weighted by the mol% of each species (WT, set 1/set 2: CE 0.86/0.94, TAG 1.92/2.29, DG 1.13/1.44, PC 1.33/1.32, PE 2.40/2.38, PS 1.61/1.62, PI 2.01/1.98 and SM 1.17/1.20 double bonds per species). Set 1: ABCA1, ANO6, ATP11A, ATP11C, ATP8A1, PLSCR1 and TMEM30A with WT, n = 2 samples each; set 2: ABCA7, ATP8B1 and ATP8B2 with WT, n = 3 samples each.

**Extended Data Fig. 4.**
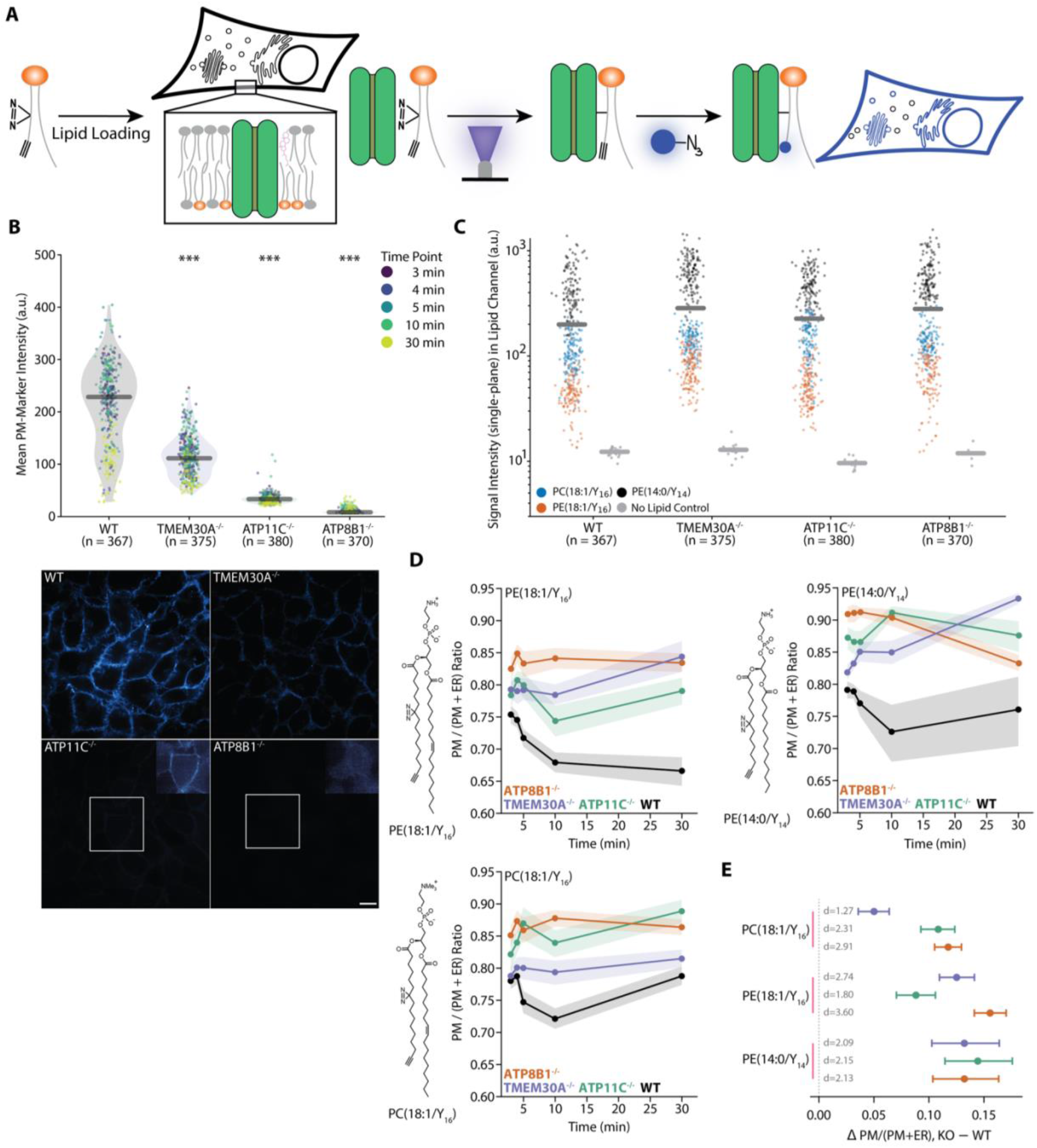
Organelle membrane partitioning of bifunctional lipids in flippase KO cells. (**A**) Schematic representation of the lipid imaging workflow. Bifunctional lipids were loaded into the plasma membrane of cells via alpha-methyl-cyclodextrin mediated lipid exchange. After a 3 min pulse, the lipid loading solution was removed and intracellular bifunctional lipids were crosslinked after defined chase time using UV light. Cells were chemically fixed immediately afterwards and covalent lipid protein conjugates fluorescently labelled using click chemistry, followed by fluorescence microscopy. (**B**) Mean plasma membrane marker intensity per image inside the segmented membrane mask for WT and the KO lines, with points coloured by chase timepoint, bars at the median and n = number of analysed images. Stars indicate two-sided Welch t-tests of each KO against WT, Holm adjusted, *** < 0.0001. Representative marker images below visualise the same effect, with insets showing contrast-enhanced regions for the two faintest lines. All images were acquired with the same microscopy settings, scale bar = 10 µm. (**C**) Single-plane lipid channel intensity per image for the three probes and the no lipid control, which defines the channel background, bars at the mean. (**D**) Plasma membrane to (plasma membrane plus ER) ratio of the exclusive organelle partition against total time, pulse plus chase, for each probe, as also shown in Extended Data Figs. 5 and 6. Shaded bands are 95% confidence intervals from 20,000 bootstrap resamples, n = 19 to 37 images per cell line and timepoint. (**E**) Time-averaged difference in this ratio between each KO and WT, taken as the area under the curve over 3 to 30 min divided by the 27 min span, with 95% confidence intervals and the standardised effect size d in pooled within-timepoint standard deviation units.

**Extended Data Fig. 5.**
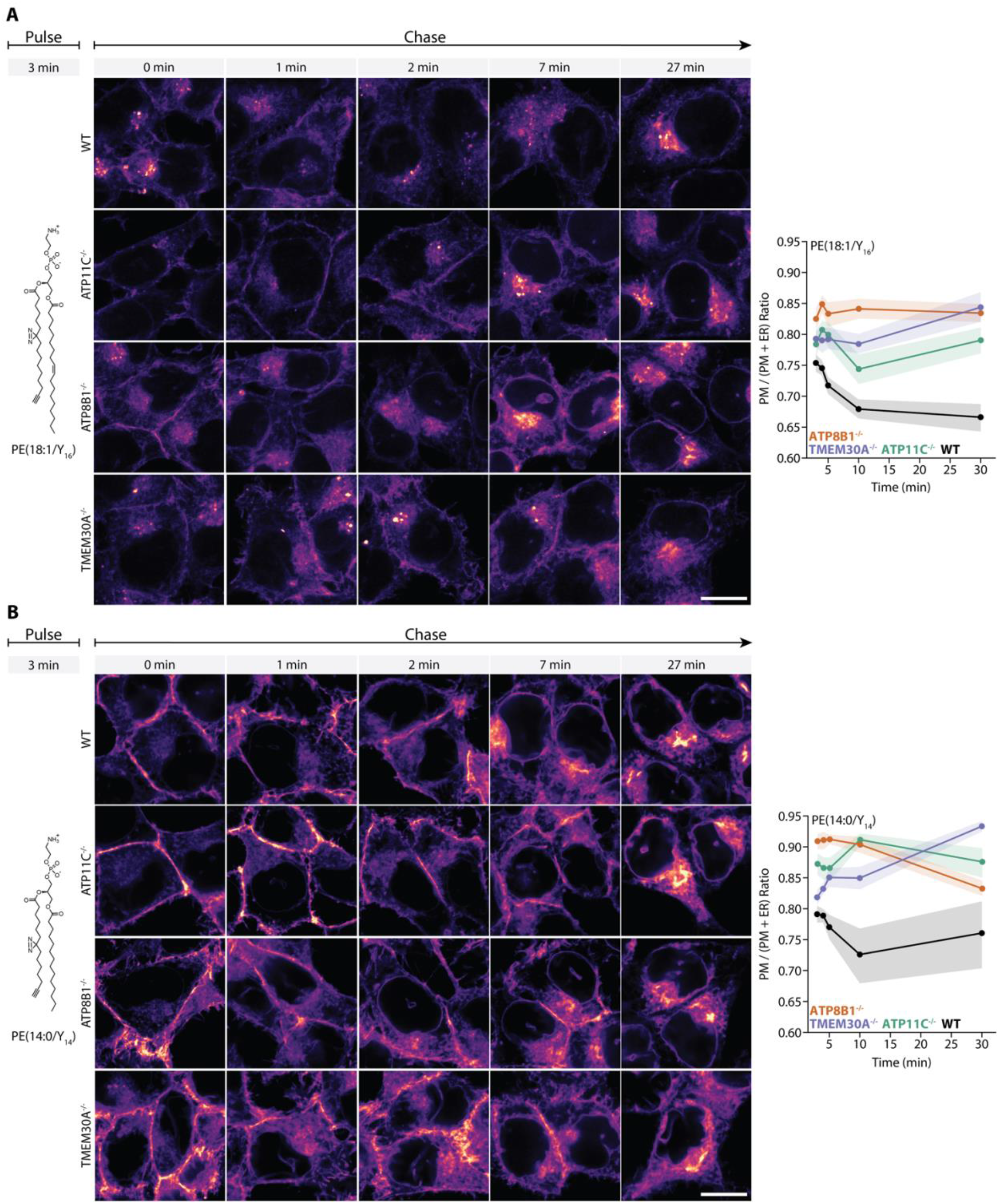
Plasma membrane retention of bifunctional PE probes in flippase KO cells. Cells were pulsed for 3 min with the indicated bifunctional lipid and chased for 0, 1, 2, 7 and 27 min before UV crosslinking, fixation and click labelling. (**A**) PE(18:1/Y_16_) and (**B**) PE(14:0/Y_14_), with the lipid structure shown on the left. Left panels, representative images of the labelled lipid in WT and KO cells at each chase time. All images were acquired with the same microscopy settings, scale bar = 10 µm. Right panels, plasma membrane to (plasma membrane plus ER) ratio of the exclusive organelle partition, calculated per image and averaged per cell line and timepoint (WT black, ATP11C^-/-^ green, ATP8B1^-/-^ orange, TMEM30A^-/-^ purple). Time is total time, pulse plus chase. Shaded bands indicate 95% confidence intervals from 20,000 bootstrap resamples, with n = 19 to 31 images per cell line and timepoint.

**Extended Data Fig. 6.**
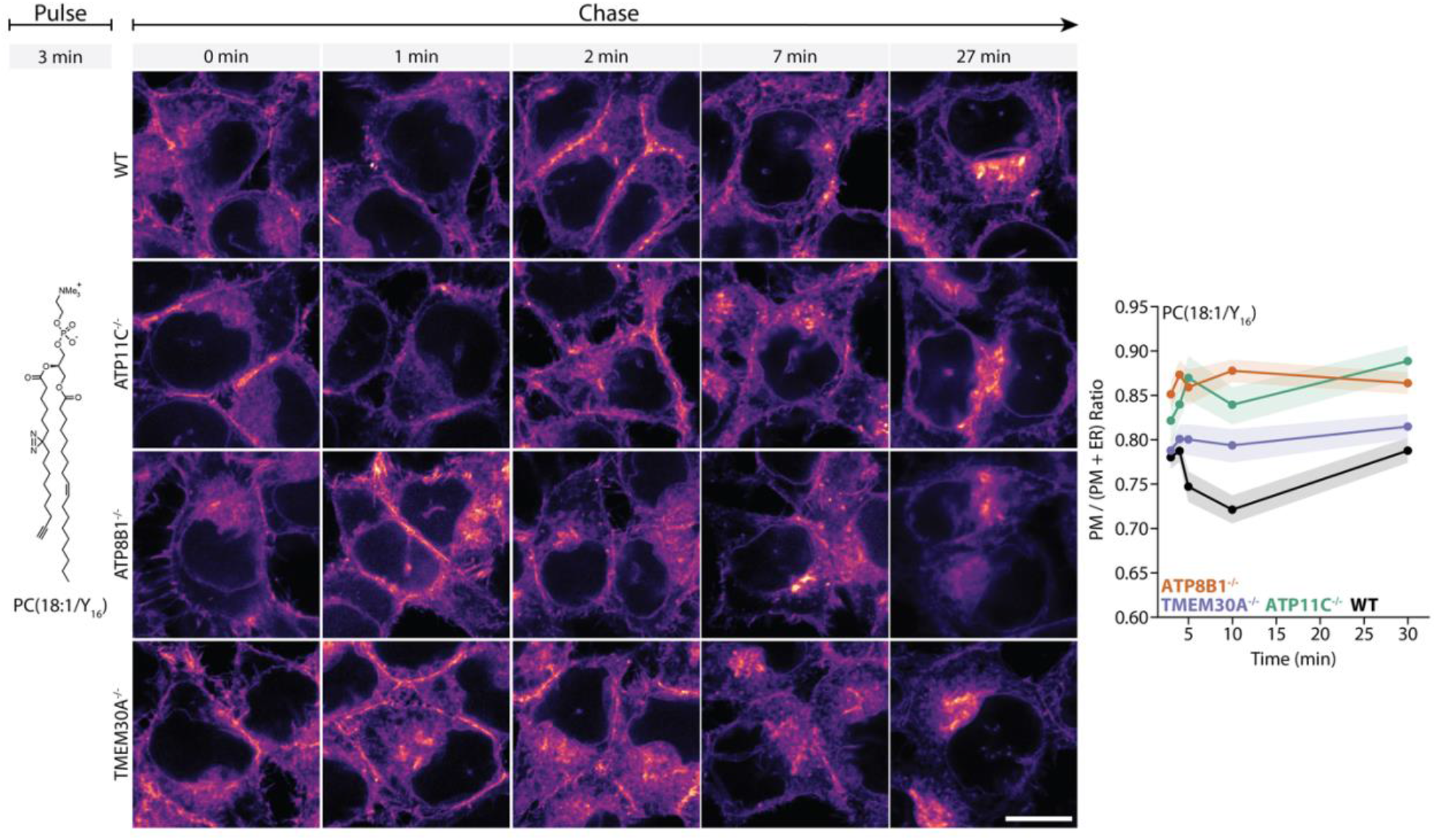
Plasma membrane retention of a bifunctional PC probe in flippase KO cells. Cells were pulsed for 3 min with PC(18:1/Y_16_), shown on the left, and chased for 0, 1, 2, 7 and 27 min before UV crosslinking, fixation and click labelling. Left, representative images of the labelled lipid in WT, ATP11C^-/-^, ATP8B1^-/-^ and TMEM30A^-/-^ cells at each chase time. All images were acquired with the same microscopy settings, scale bar = 10 µm. Right, plasma membrane to (plasma membrane plus ER) ratio of the exclusive organelle partition, calculated per image and averaged per cell line and timepoint (WT black, ATP11C^-/-^ green, ATP8B1^-/-^ orange, TMEM30A^-/-^ purple). Time is total time, pulse plus chase. Shaded bands indicate 95% confidence intervals from 20,000 bootstrap resamples, with n = 20 to 37 images per cell line and timepoint.

**Extended Data Fig. 7.**
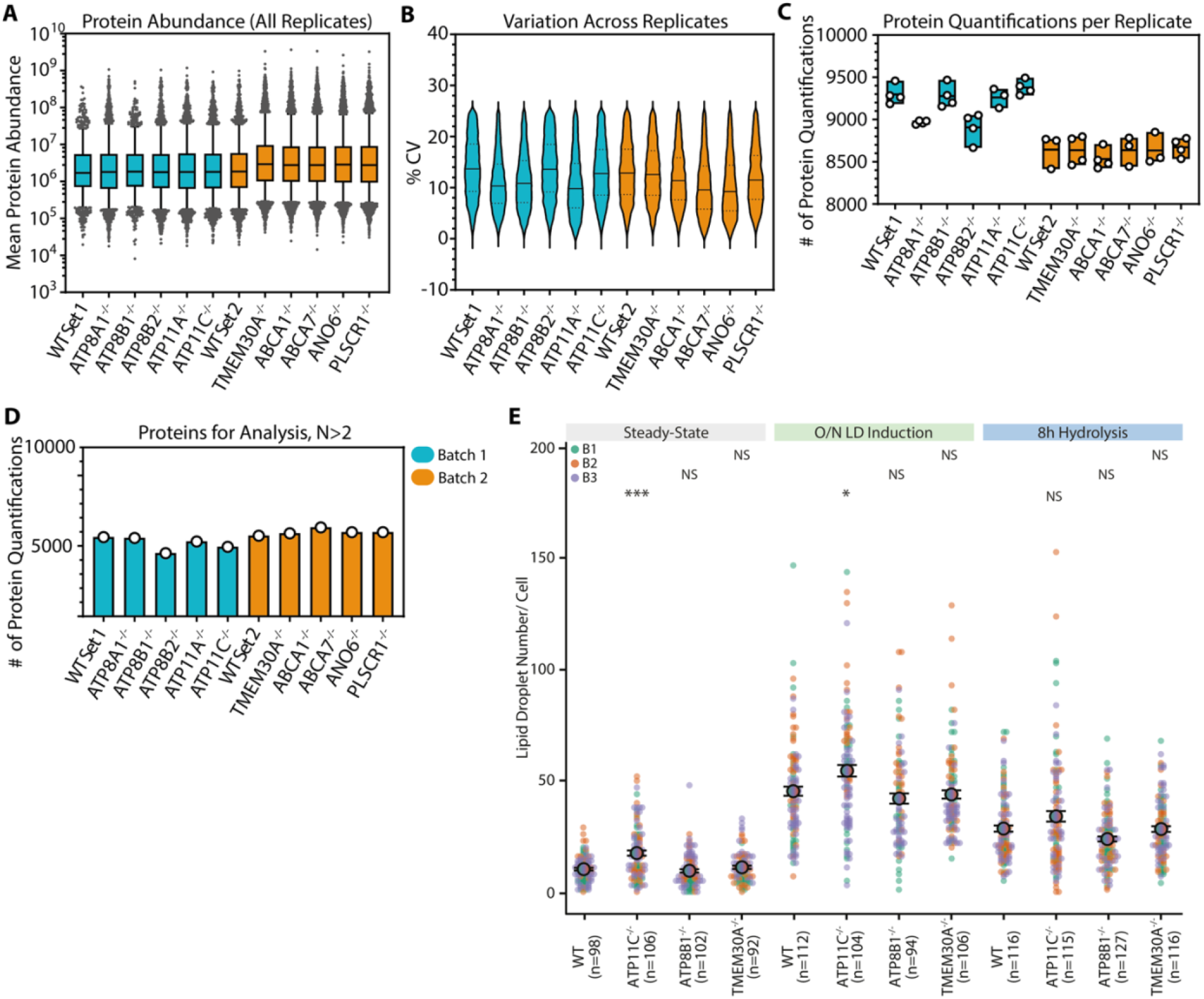
MS detection and quantification representations for proteomics datasets and lipid droplet numbers. **(A)** Mean peptide intensity for proteins detected and quantified by DIA LC-MS/MS, prior to %CV filtering (FDR < 1% for all included peptide values). Data is depicted as box-and-whiskers with whiskers spanning the 10th to 90th percentiles, line at median, and dots for remaining protein values. **(B)** Coefficient of variation (%CV) across all replicates for all proteomics data included in this study (N=4 biological replicates total, gathered in two separate batches for each CRISPR line). Shown as violin plots for the full range of data, with solid line at the median and dotted lines at quartiles. **(C)** Number of protein quantifications per replicate for each CRISPR cell line, prior to %CV or replicate completeness filtering (FDR <1%). Each dot indicates a replicate and the box shows the min/max with solid line at the median. **(D)** Protein abundance values used for all analyses in all main and supplementary figures in this study were filtered for CV < 25% and data completeness in N>2 biological replicates. (E) Average LD number per cell for the oleic acid feeding experiment shown in Fig. 4C. Data points are colored by biological replicate (B1-B3), with n = number of analyzed cells, p-values derived from a two-sided Mann-Whitney-U test for each KO compared to WT, and error bars indicating standard error of the mean. p-Values: *** < 0.0001, ** < 0.001, * < 0.01, NS = not significant.

**Extended Data Fig. 8.**
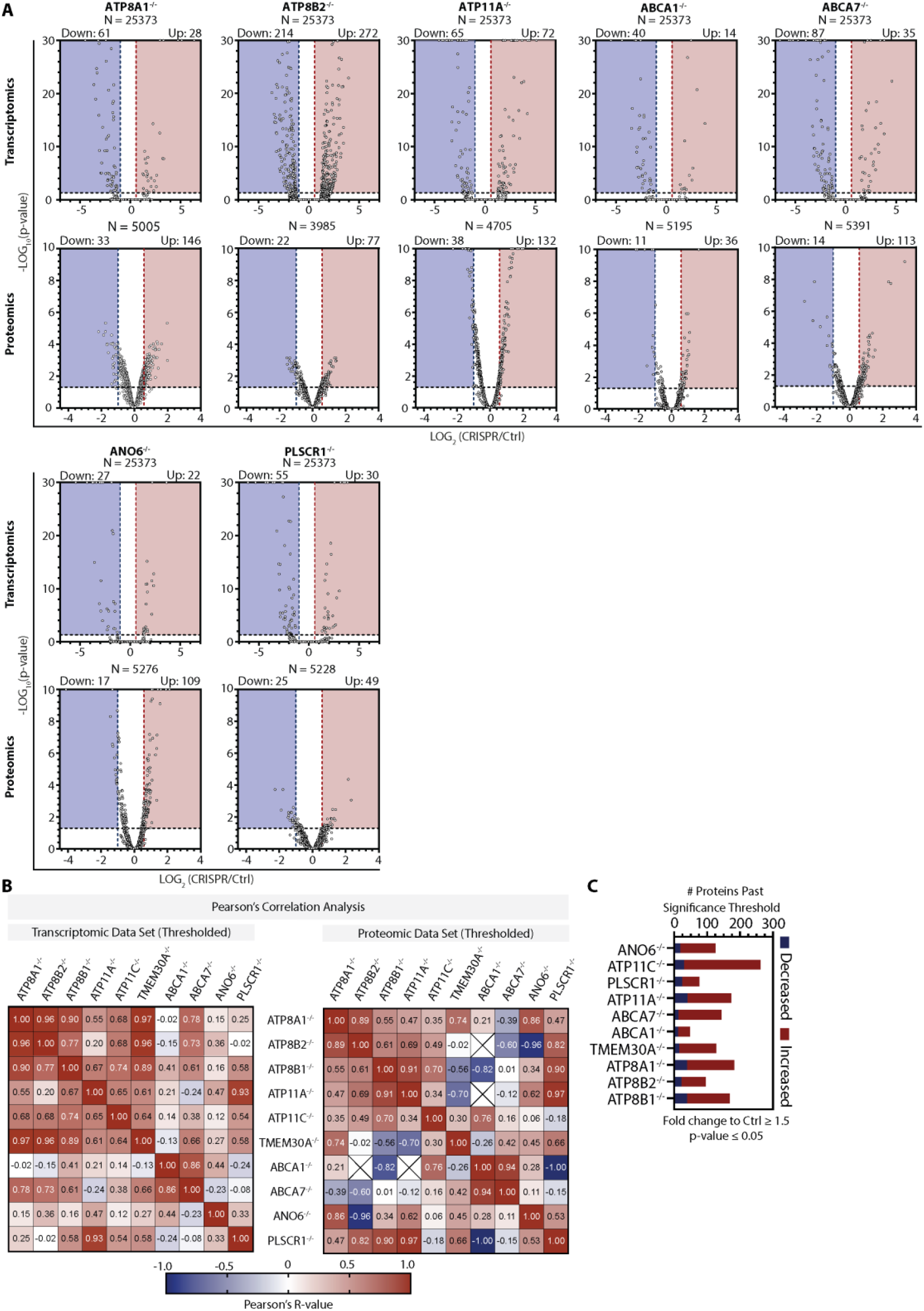
Transcript and protein abundances are differentially modulated by CRISPR KO of plasma membrane lipid asymmetry-associated proteins. **(A)** Volcano plots showing the full data distribution for transcriptomics (top panels) and proteomics (lower panels) analyses in CRISPR knockout lines (genes indicated above graphs; ATP8B1, ATP11C, and TMEM30A volcano plots are in Figure 3). Shaded regions represent fold-change to wildtype ≥2 (x-axis) and p < 0.05 (y-axis), and each dot represents a transcript/protein, with total counts indicated above each graph. Data included here is already filtered for transcriptomic and proteomic quality as described in Methods. **(B)** Data correlation matrices showing Pearson’s Correlation R values for each pair of CRISPR KOs in the transcriptomic (left) and proteomic (right) data for significantly regulated gene products only (thresholded by +/ −50% change to control, p < 0.05; N = 867 transcripts and N = 1037 proteins). R values for each comparison are written in each cell. **(C)** Number of proteins passing significance thresholds for each CRISPR condition. For all CRISPR cell lines, the number of significantly increased protein abundances (red bars) exceeded the number decreased (blue bars), and the total (sum of stacked bars) varied across CRISPR conditions.

**Extended Data Fig. 9.**
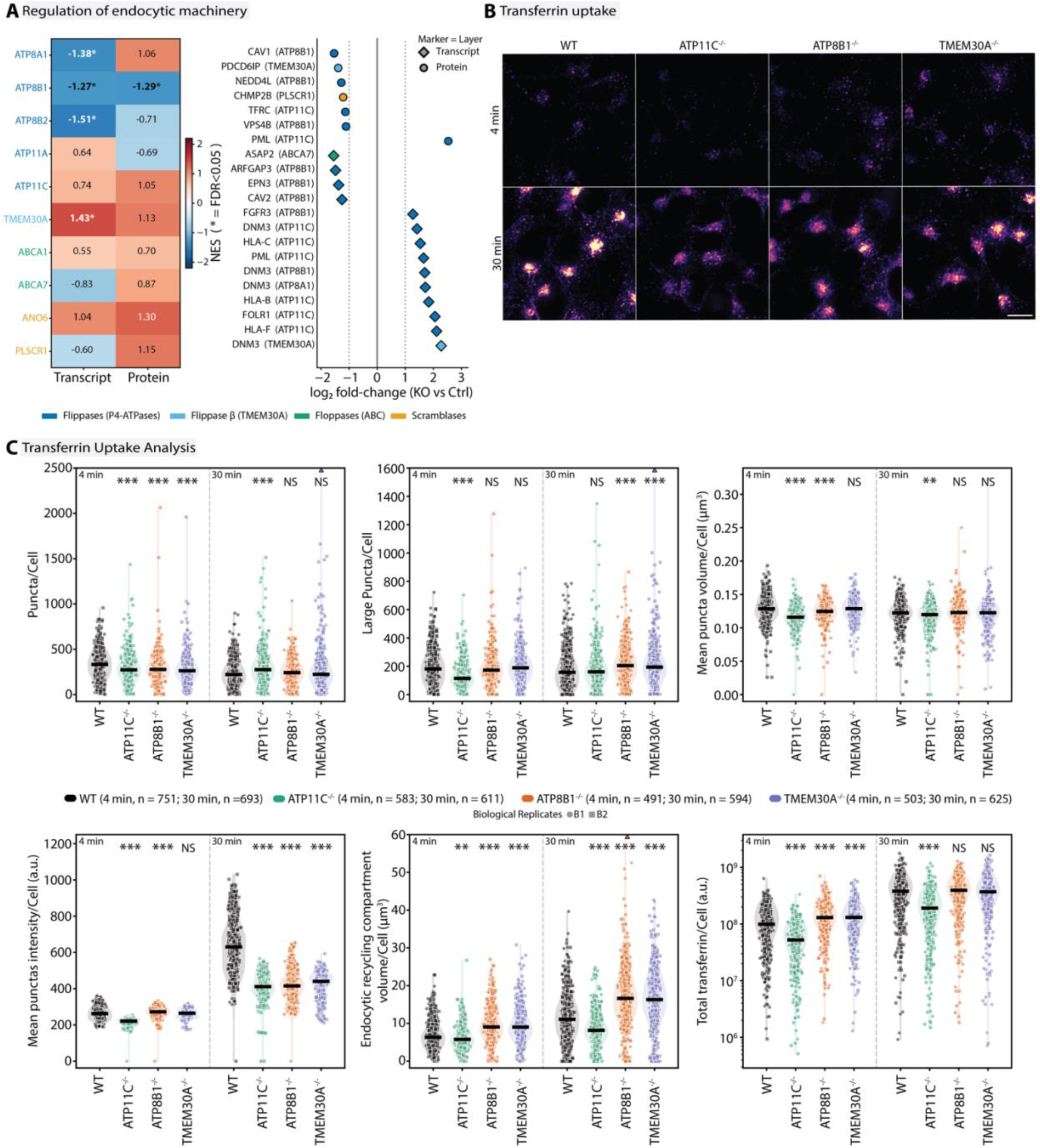
Lipid translocase KOs affect the endocytic machinery and alter transferrin uptake in a gene-specific manner. **(A)** Left: normalised enrichment score (NES) of the endocytosis-associated gene set per KO line, for transcript and protein; asterisks indicate FDR < 0.05. Right: regulated endocytic gene products per KO line as log2 fold - change to control; shape denotes the layer (diamond, transcript; circle, protein), colour the transporter class, dotted lines a two-fold change. **(B)** Alexa Fluor 647 transferrin in WT and the ATP11C^-/-^, ATP8B1^-/-^ and TMEM30A^-/-^ lines after 4 and 30 min of uptake. All images were acquired with identical settings, scale bar = 10 µm. **(C)** Per-cell puncta number (5 to 500 voxels), large puncta (above 500 voxels), mean puncta volume, mean puncta intensity, proposed endocytic recycling compartment volume (largest transferrin-positive structure) and total transferrin. Points are single cell measurements coloured by cell line, shape indicating biological replicate (B1, B2), line at the median. n = number of analyzed cells. p-Values from a two-sided Mann-Whitney-U test for each KO compared to WT at the matching uptake time, Benjamini-Hochberg corrected; *** < 0.001, ** < 0.01, * < 0.05, NS = not significant. Points above the axis range are drawn as triangles; zeros are absent from the log axis.

**Extended Data Fig. 10.**
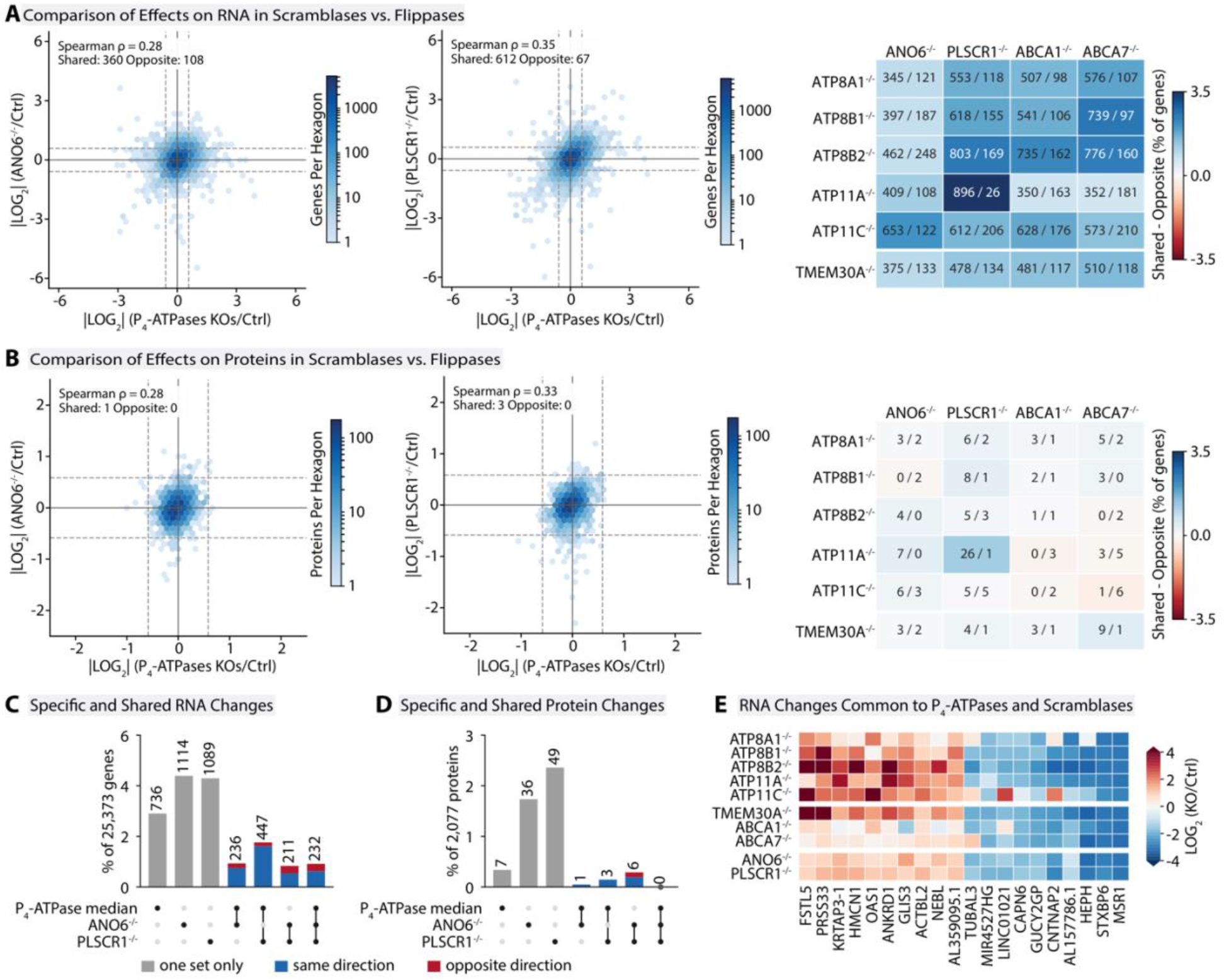
Comparison of transcript and protein abundance responses across lipid translocase knockouts. **(A)** Transcriptomic comparisons. Left panel, gene-wise median log_2_ fold-change relative to control across five P_4_-ATPase KOs (ATP8A1, ATP8B1, ATP8B2, ATP11A and ATP11C; horizontal axis) compared with ANO6 KO (vertical axis). Middle panel, the same P_4_ median compared with PLSCR1 KO. Right panel, shared and opposite responses for each individual P_4_-ATPase KO and TMEM30A KO compared with ANO6, PLSCR1, ABCA1 and ABCA7 KOs. **(B)** Corresponding proteomic comparisons with ANO6 (left panel), PLSCR1 (middle panel) and individual KO pairs (right panel). TMEM30A was analyzed separately and was not included in the P_4_ median. In the left and middle panels, hexagon color indicates the number of genes or proteins on a logarithmic scale, with density scales matched within each assay. Spearman correlation coefficients are indicated. Solid lines mark zero change and dashed lines mark reciprocal 1.5-fold thresholds (log_2_ fold-change ≥ log_2_(1.5) or ≤ -log_2_(1.5)). Shared and opposite counts require both compared values to reach these thresholds in the same or opposite directions, respectively; for the P_4_ median, this requires at least three of five KOs. In the right panels, cell labels give shared/opposite counts and color indicates 100 × (shared - opposite)/N, with a common scale across A and B. Blue indicates more shared responses and red more opposite responses. Analyses used fixed sets with unique human Ensembl identifiers and available fold-changes in all ten KOs (N = 25,373 RNA genes and 2,077 proteins represented by unique gene identifiers), without significance filtering or imputation. **(C)** Transcripts changed at least 1.5-fold in the P4 median only, in ANO6 KO only or in PLSCR1 KO only (grey), or in two or all three of these (dots below the bars), shown as percentage of the 25,373 genes with counts above the bars; blue, same direction; red, opposite direction. **(D)** The same for the 2,077 proteins. **(E)** Log2 fold-change relative to control in all ten KO lines for the 20 transcripts with the largest P_4_ median change among the 157 that changed in the same direction in the P_4_ median and in both ANO6 and PLSCR1 KOs; red, increase; blue, decrease.

**Extended Data Fig. 11.**
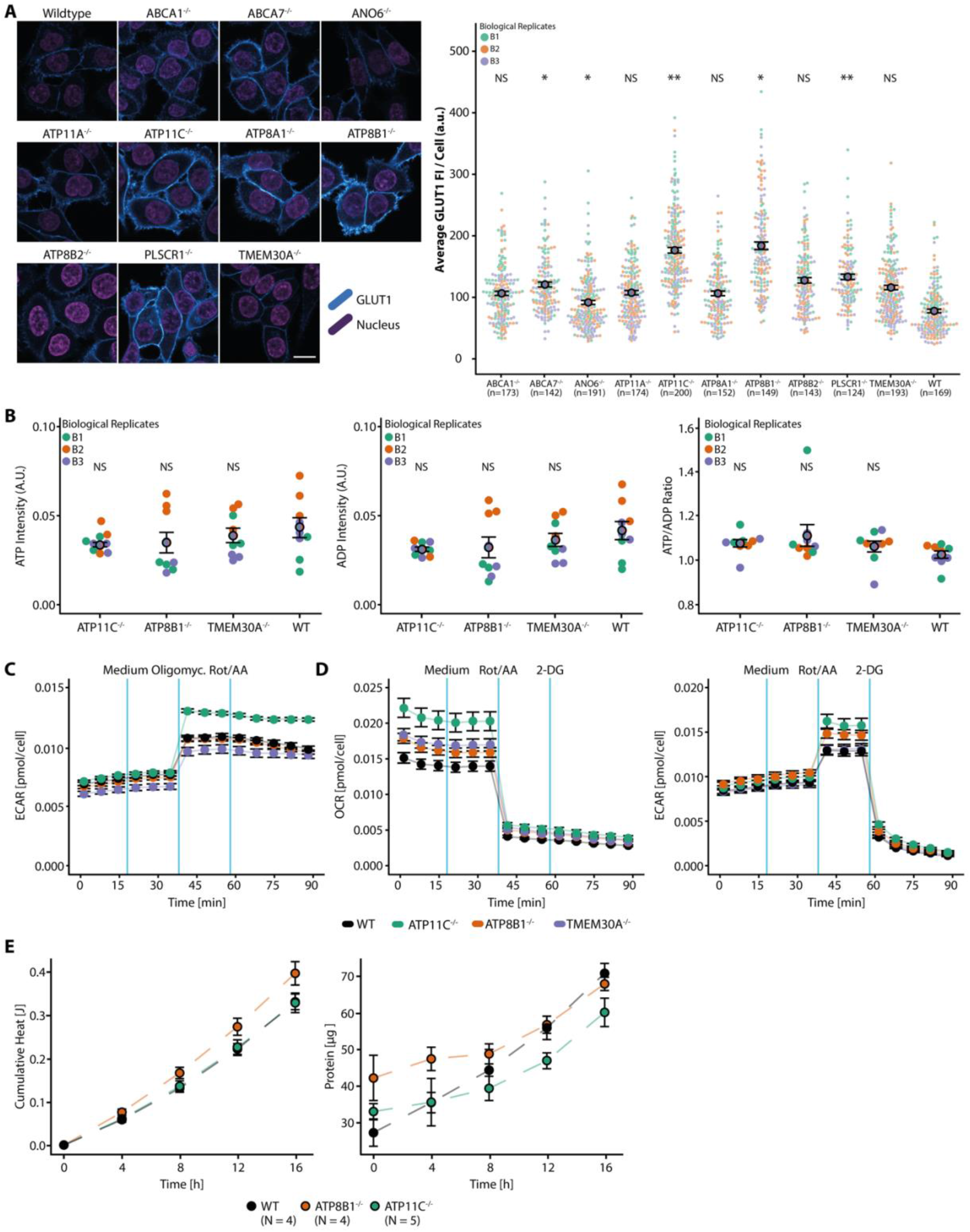
KO cell lines exhibit altered bioenergetics. **(A)** Immunofluorescence staining of GLUT1 in all KO cell lines (left panel) and analysis of mean fluorescence intensity (FI) per cell (right panel). All images were acquired with the same microscopy settings, scale bar = 15 µm. Images were analysed with Fiji and average fluorescence intensities per cell were determined. Coloured points show single cell measurements with colours indicating biological replicate, black circles show averages and error bars indicate standard error of the mean. Statistical analysis was performed with Student’s t-test where all KOs were tested against the WT. **(B)** ATP and ADP levels (left and middle panels) and ATP:ADP ratio (right panel) in selected KO lines as determined by mass spectrometry. Error bars indicate standard error of the mean. Colored dots represent three biological replicates (B1-B3). **(C)** Seahorse assay to assess extracellular acidification rates (ECAR) after addition of Oligomycin and Rotenone (Rot)/Antimycin A (AA). Experiment was repeated 3 times and blue lines indicate timing of medium/inhibitor addition. Data is from the same experiment as shown in Fig. 5D. Error bars indicate standard error of the mean. **(D)** Oxygen consumption rates (OCR) and ECAR after the addition of Rot/AA and 2-Deoxy-D-Glucose (2-DG) (data from an independent experiment). Experiment was repeated 3 times and blue lines indicate timing of medium/inhibitor addition. Error bars indicate standard error of the mean. **(E)** Cumulative heat measured by nano-calorimetry over time (left panel). Protein amount in nano-calorimetry samples over time (right panel). N indicates the number of biological replicates. Error bars indicate standard error of the mean. Dashed lines are linear interpolations. p-Values: *** < 0.0001, ** < 0.001, * < 0.01, NS = not significant.

