## Supplementary Information for "Disruption of Plasma Membrane Lipid Asymmetry Alters Cellular Energetics"

### Materials and Methods

Cell Culture

Cells were cultured in McCoy's 5A GlutaMAX™ medium (36600-021, Life Technologies) supplemented with 10 % fetal bovine serum (10270-106, Life Technologies, not-heat inactivated) and 100 µg/ml antibiotic Penicillin-Streptomycin (10 000 U/mL, 15140-122, Life Technologies). Cells were trypsinised with 0.05 % trypsin (25300054, Life Technologies) and passaged every 48 h to 72 h in a 1:10 ratio (ATP11C$^{-/-}$ in a 1:3 ratio) into Nunc EasYFlask 75 cm2 Nucleon Delta Surface (156499, ThermoFisher Scientific) flasks. Cells were grown under standard cell culture conditions (37 °C, 5 % $CO_2$). For all experiments, cells were used to maximal passage 10 after thawing.

Knockout Generation using CRISPR/Cas9

Knockout lines were generated by deletion of critical exons (exon present for all isoforms of one protein) from endogenous genes. For each target region, the sequence was analyzed for appropriate guide RNA binding sites using the Geneious software and the CRISPOR design webtool (http://crispor.tefor.net/). Four combinations of guideRNA pairs were tested in advance, and the pair with the best performance was used to generate the knockout clones. Cuts were made using guide RNA pairs flanking the critical exon. Cas9 protein, guideRNAs (ordered as crRNA) and tracr-scaffoldRNA were obtained from IDT (Integrated DNA Technologies). The following guideRNAs were used for the respective genes:

**Table 1 | Overview of used guideRNAs for KO generation**

| Target Gene | 5'-Guide | 3'-Guide |
|---|---|---|
| ABCA1 | GGGGTTCATGATACCAGCGTTGG | ATCCATGTAAGTATCAGATCAGG |
| ABCA7 | TCCACCCCCATCGGGTCGCCCGG | TATGATAGTATGGTCTGCCTGGG |
| ANO6 | CGATTGGGTGTTACAATCCTGGG | GTGCTGAGCTTCGTTGGCCAGGG |
| ATP11A | GTCAACACAAACCCAGACGCTGG | ACGGACAAGGGTTTCCACGGCGG |
| ATP11C | AAGATAACCTGCCGCAAGGAGGG | AACCTACTAATTGCATGAAGAGG |
| ATP8A1 | AAGAAGTTGAACACTTACGGGGG | TATCTATGCAGCATCGGCAGAGG |
| ATP8B1 | ACATCTTGATGCCTCGAGGAAGG | TGATGATTATCAGTAGCCCCAGG |
| ATP8B2 | TACCGCGACTAGATCTGCAGTGG | GGCAAGGACAACCTACCGTTGGG |
| PLSCR1 | GTATCTGATCTATCTATAGCAGG | AGTTAGTGTCAATATCCCGAGGG |
| TMEM30A[1] | GTTAGCATCGACTTTTCACAGGG | GAGATTTCACGTAACGACGATGG |

HCT116 cells were transfected with Cas9 ribonucleoprotein complexes using Neon electroporation kit and device (Invitrogen). For the preparation of the crRNA-tracrRNA duplexes, RNA oligos (crRNA and tracrRNA) were mixed in equimolar amounts (100 µM). The mixes were incubated for three minutes at 95 ˚C and afterwards slowly cooled to RT. In order to form the Cas9 ribonucleoprotein complexes, crRNA-tracrRNA duplexes were mixed with Cas9 in a 1.4:1 molar ratio. The mixes were incubated for 15 min at RT. Subsequently, the electroporation mixture was prepared by combining Cas9 ribonucleoprotein and cell suspension in electroporation buffer.

From this mixture, 10 µL were used for electroporation that was performed with the following settings: 1530 V, 20 ms pulse width, 1 pulse. After 72 h cells were harvested and sorted as single cells into 96-well plates (167008, Thermo Fisher Scientific) by FACS. Single-cell clones were screened for exon deletion by PCR using primers spanning the targeting exon. To verify the absence of the wild-type allele in homozygous knockout candidates a flanking PCR with one primer upstream of the deletion and one primer inside the deleted sequence was performed. Deletions were identified by Sanger Sequencing.

RNA Sequencing and Analysis

RNA was isolated from the respective cell lines using RNAeasy Mini Kit (74104, Qiagen) and RNA quality was assessed by RNA integrity number (RIN). RIN was determined with the Bioanalyzer 2100 (Agilent). Only RNA with a RIN >9 was used for sequencing. RNA was enriched by ribosomal RNA depletion (RNaseH) using the NEBNext rRNA Depletion Kit (E6310, New England Biolab). Libraries were prepared using the NEBNext Ultra II Directional RNA Library Prep Kit for Illumina (E7760, New England Biolabs) and sequenced with NextSeq 500 (Illumina). Three biological replicates were prepared and sequenced.

30 single-end and 24 paired-end Illumina read data sets of sizes between 37.6 Mio and 138.8 Mio (median 43.9 Mio) were processed. Illumina universal adapters were trimmed from both ends of the reads with Cutadapt v1.16; discarding reads shorter than 15 nt led to read data sets of sizes between 37.5 Mio and 138.8 Mio (median 43.9 Mio). Quality of reads was assessed using FastQC 0.11.9. Reads were mapped against the Homo sapiens genome reference assembly GRCh38 and genes of the Ensembl release v99[2] were quantified using STAR 2.7.3a (settings: --outSAMstrandField intronMotif --outFilterMultimapNmax 1 --outSJfilterCountUniqueMin 3 1 1 1 --quantMode GeneCounts --alignSJDBoverhangMin 5 --alignSJoverhangMin 8 --outFilterMultimapNmax 1 --outFilterMismatchNmax 999 --outFilterMismatchNoverLmax 0.1 --alignMatesGapMax 200000 --seedSearchStartLmax 25 --chimSegmentMin 20 --twopassMode Basic --alignIntronMin 20 --alignIntronMax 200000).[3] Read coverage plots for genetic loci removed by CRISPR/Cas9 were plotted with R package Gviz 1.34.1. Genes with at least 10 reads in at least one sample were input into the analysis of differential gene expression with DESeq2 1.22.1.[4] Three samples (1 x WT, 1 x PLSCR1, 1 x ATP8A1) were identified as outliers and discarded as their positions were far from the other replicates in PCA. Sets of differentially expressed genes are those with an FDR<1% considering p-values from the Wald test implemented in DESeq2 testing for a fold-change of at least two-fold up or down.

For transcriptomic data presentation in all main figures, Extended Data Fig. 8A, and Supplementary Table 3, the complete dataset of quantified gene abundances are included (no additional filtering). K-means clustering of gene abundance fold-changes was performed by Morpheus (software.broadinstitute.org/Morpheus), using Euclidean distance at maximum iterations = 1000. Gene ontology enrichment tests were performed for Clusters 4 (up) and 5 (down) with PantherDB[5,6] (pantherdb.org) against the total human genome, using a Fisher's Exact test for statistical over-representation. For the correlation plot in Extended Data Fig. 8B, only genes that exhibited a significant up- or down-regulation (threshold = fold-change to wildtype control of +/- 50% and p-value < 0.05) in at least one KO were included, using the Pearson Correlation test for XY matrices (two-tailed, 95% confidence interval). For the functional classification graphs in Figure 4A and Figure 5A, abundances for all genes mapping to the indicated functional pathways are plotted (diamond symbols) while only those passing significance thresholds (+/-50% and $P < 0.05$ to control) are highlighted by solid color. All gene ontology terms and mapped genes/proteins thereof are included in Supplementary Tables 5 and 6, with abundance and p-values included. All heatmaps and graphs were generated using GraphPad Prism. The complete transcriptomics dataset is available as an Excel matrix in Supplementary Table 3. Transcriptomics raw files were submitted to the Gene Expression Omnibus[7] (GEO) repository with the identifier GSE276056 (https://www.ncbi.nlm.nih.gov/geo/). For the comparison of lipid translocase KOs (Extended Data Fig. 10A,B), log2(KO/WT) values of the 25,373 genes and 2,077 proteins quantified in all ten KO lines (Supplementary Tables 3 and 4) were analyzed in Python without significance filtering. The gene-wise median of the five $P_4$-ATPase KOs was compared with $ANO6^{-/-}$ and $PLSCR1^{-/-}$ by Spearman rank correlation. Shared and opposite responses were counted for this median and for each $P_4$-ATPase KO and $TMEM30A^{-/-}$ against $ANO6^{-/-}$, $PLSCR1^{-/-}$, $ABCA1^{-/-}$ and $ABCA7^{-/-}$, using reciprocal 1.5-fold thresholds. For Extended Data Fig. 10C to E, each gene or protein was assigned to the combination of the P4-ATPase median, $ANO6^{-/-}$ and $PLSCR1^{-/-}$ in which it passed these thresholds, and a combination was counted as same direction when all changes had the same sign.

Lipidomics Sample Preparation, Measurements and Analysis

HCT116 cells were seeded two days prior to lipid extraction with $1 \times 10^6$ cells in Nunclon™ Delta Surface dishes (150326, Thermo Scientific). Cells were washed twice with ice-cold PBS and subsequently scraped (3010, Corning) in 500 µL ml IPA/$H_2O$ (1:1, v/v) and suspended using 0.5 mm stainless steel beads and TissueLyser II (Qiagen). An aliquot of 3e5 cells was transferred to a fresh Eppendorf tube and dried under vacuum with a Speedvac. For lipid extraction, 700 µL of internal standard mixture in MTBE/MeOH (10:3 v/v) and 140 µL water were added to the residue.[8] The suspension was mixed for 1.5 h at 4 C, followed by 30 min centrifugation at 13000 x g at 4 C. Before the analysis, 50 µL lipid extract were transferred to a 96well plate, dried under vacuum and resolubilized in 100 µL spray solution containing isopropanol/methanol/chloroform (4:2:1 v/v/v) with 7.5 mM ammonium formate. The diluted lipid extracts were subjected to nanoelectrospray ionizations using a robotic ion source Triversa Nanomate (Advion).[9]

FTMS and FTMSMS raw spectra and transient data were acquired with a QExactive Orbitrap FTMS instrument (Thermo Fisher Scientific) and a connected Booster X2 system (SpectroSwiss). On the QExactive, slens level was set to 10% and capillary temperature to 275C. Spectra were acquired in two polarity modes by two separate methods:

1) Ultra-high resolution FTMS using tSIM acquisition of 40 Th windows with 20 Th steps. For UHR, 2 s transients were recorded to achieve 1M resolution at m/z 200.

2) For FTMSMS, regular 0.5 s transients were recorded to give 280k resolution fragment spectra. All mass bins in the range of m/z 400 - 1000 were fragmented in 1 Th steps at NCE 15% and 22%, respectively.

Post-acquisition, transient data were processed to spectra by PeakbyPeak software (SpectroSwiss) using aFT mode. Intensities of the spectra were ion injection time normalized, transients with the same scan header were averaged and spectra were exported to mzML file format. tSIM spectra were further subjected to Simtrim application to cut the isolation biased m/z ranges. Lipid identification was performed by Lipidxplorer 1.2.4.[10] Chol, CE, TG, were analyzed via UHR FTMS+, DG from FTMSMS+, Cer, HexCer, SM, PC, PE, PI, CL and corresponding lysospecies via FTMS-, and PS, LPS via FTMSMS-. Obtained lipid class and species data was plotted in R and with Omicsvolcano[11]. The complete lipidomics dataset is available as an Excel matrix in Supplementary Table 7.

Proteomics Sample Preparation, Measurements and Analysis

Cells were seeded two days prior to proteomics sample collection with 5 x $10^5$ cells in 60 mm Nunclon™ Delta Surface dishes (150326, Thermo Scientific). Cells were washed twice with ice-cold PBS and subsequently scraped (3010, Corning) in 500 µL lysis buffer (RIPA buffer with EDTA and EDGA (J60645, Alfa Aesar), 2x protease inhibitory cocktail (11697498001, Roche), 5 % SDS (151-21-3, Serva) and 1 % n-octyl-beta-D-Glucopyranoside (O8001, Sigma-Aldrich)) and snap-frozen in protein LoBind Eppendorf tubes (90410, Thermo Scientific). Samples were stored at - 80°C until further processing in 2 batches (wild-type included in each batch) in a block-randomized fashion, 4 replicates per condition. Samples were lysed with the addition of 3 micro spatulas of 0.5 mm steel beads (Next Advance, Inc.) in a TissueLyser II (QIAGEN) operated at 30 Hz for 15 min at 4 °C. The lysate was centrifuged for 10 min at 13000 x g at RT, vortexed for 5 s, and centrifuged again as before. Proteins in the supernatant were precipitated at a final concentration of 85% acetone and 30 mM NaCl at RT for 30 min under agitation.[12] Precipitated protein pellets were resuspended in 1x S-trap lysis buffer and the protein concentration was determined using Pierce BCA protein assay kit (10678484, Thermo Scientific). Per sample, 45 µg protein were digested overnight using S-traps (ProtiFi, LLC) and Trypsin/Lys-C Mix, Mass Spec Grade (Promega) at a ratio of 0.7 µg enzyme per 10 µg sample protein. The resulting peptide amounts were measured using the Pierce quantitative colorimetric peptide assay (23275, Thermo Scientific) and 0.15 µg/µl in 0.2% formic acid (FA, Merck KGaA) were used for LC-MS analysis.

The LC-MS setup (all Thermo Scientific/ PharmaFluidics, exceptions noted) consisted of an UltiMate 3000 UHPLC system equipped with an Acclaim PepMap precolumn (100 µm x 20 mm, C18, 5 µm, 100 Å) and a 50 cm µPAC pillar array column coupled via µPAC Flex iON interface plus nESI emitter (20 µm, 5 cm, Fossiliontech) to a Q Exactive HF. 5 µl sample were loaded at a flow rate of 5 µl/min and eluted using a 2-sloped linear gradient (all steps 0.5 µl/min, 0.1 % FA) from 0 - 17.5 % ACN in 100 min and 17.5 - 35 % ACN in 50 min. With interspaced 22 min gradient blanks this results in a turnover of 6 samples per day.

Data-independent acquisition (DIA) consisted of a Full MS scan (60k, 3e6 AGC, 40 ms IT, centroid) and a series of 32 MS2 scans (30k, 1e6 AGC, 55 ms IT, 18 m/z window, 24 NCE, first mass 100 m/z, centroid) covering 400 – 966 m/z in a staggered DIA fashion[13] at a rate of ca. 3.9 scans per chrom. FWHM. Raw files were demultiplexed and converted to mzML with MSConvert and processed using DIA-NN 1.8.[14] The MaxQuant contaminant fasta and a human database (Uniprot SWISS-PROT reference proteome UP000005640 canonical+isoform, 24.11.2021) were used to generate an in-silico spectral library (20501 proteins, and 20298 genes) for searching mzML files with the following settings: --cut K*,R* --var-mods 1 --var-mod UniMod:35,15.994915,M --use-quant --double-search --individual-mass-acc --individual-windows --smart-profiling --pg-level 2 --species-genes --peak-center --no-ifs-removal --report-lib-info --il-eq --matrix-qvalue 0.01 --nn-single-seq --fixed-mod UniMod:39,45.987721,C --strip-unknown-mods.

Using DIA-NNs protein group (pg) matrix at 1% FDR, entries were processed with an in-house R script. In short, entries were considered for analysis if the data completeness was at least 50 %

and the coefficient of variation (CV) is below 25 % in both a particular experimental condition and the respective wild-type from the same preparation batch. The MS detection and quantitation variability is depicted in Extended Data Fig. 7A-D. For all other figures containing proteomics data, differential expression analysis was performed using limma 3.50.0 with $\alpha = 0.01$ and $|\log2 \text{ fold-change}| > 0.5$.[15] The complete proteomics dataset is available as an Excel matrix in Supplementary Table 4. Proteomics raw and intermediate files were submitted to the MassIVE[16] repository with the identifier MSV000089574 (https://massive.ucsd.edu/ProteoSAFe/static/massive.jsp).

Proteomics data analysis & representation

For k-means clustering and gene ontology analysis of protein abundances across all CRISPR samples (Figure 3, Extended Data Fig. 8), k-means clustering and gene ontology analyses were performed using the same parameters as in the transcriptomics dataset analyzed in parallel (Morpheus and PantherDB; see Transcriptomics Methods above). For Figure 3A and Extended Data Fig. 8, all protein abundances passing DIA-MS data quality filters are shown (N=7118 proteins across all CRISPR cell lines), and gene ontology analysis was only performed for the clusters exhibiting average up (Cluster 4) or down (Cluster 5) trends. For Figure 3B, the "thresholded" proteomic data was filtered for protein abundances with a fold-change to control of +/- 50% and p-value < 0.05 in at least one CRISPR condition (N=1067 proteins). Data correlation analysis (Extended Data Fig. 8B) was also performed on the thresholded data (N=1067 proteins) in GraphPad Prism, using the Pearson Correlation test for XY matrices (two-tailed, 95% confidence interval). For the volcano plots in Figure 3C and Extended Data Fig. 8A, all MS quality filtered data is shown for each CRISPR condition (N>3900 proteins quantified in all replicates). For the functional classification graphs in Figure 4A and Figure 5A, abundances for all proteins mapping to the indicated functional pathways are plotted (circle symbols) while only those passing significance thresholds (+/-50% and P < 0.05 to control) are highlighted by solid color. All gene ontology terms and mapped genes/proteins thereof are included in Supplementary Tables 5 and 6, with abundance and p-values included. All heatmaps and graphs were assembled in GraphPad Prism.

Protein comparisons (Extended Data Fig. 10B) were performed in the same way with the log2(KO/WT) values of the 2,077 proteins quantified in all ten KO lines (Supplementary Table 4).

Imaging and Quantification of Plasma Membrane Phosphatidylserine Exposure

HCT116 WT cells and the ten knockout lines were seeded in glass bottom multiwell plates two days prior to the experiment with 4000 cells per well. The outer leaflet of the plasma membrane was labelled with the recombinant phosphatidylserine biosensor LactC2-eGFP[17], applied extracellularly at a final concentration of 0.5 µM for 10 min at 37 °C, and the plasma membrane was counterstained with CellMask Deep Red. Live cells were imaged with identical settings on an Olympus IX83 with a Yokogawa CSU spinning disc (excitation 488 nm for eGFP and 640 nm for CellMask Deep Red, lateral pixel size 0.043 µm), and three fields from one well were acquired per cell line.

Images were analyzed in Python. Both channels were median filtered, corrected for the camera offset and background subtracted with a rolling ball (50 pixel radius). The plasma membrane was segmented in the CellMask channel by Otsu thresholding, and bright compact objects and intense puncta in the biosensor channel were excluded from the analysis. PS exposure was quantified per field as the mean LactC2-eGFP intensity within the plasma membrane mask.

Cyclodextrin Extraction and Lipidomic Analysis

HCT116 WT cells and the TMEM30A and ATP11C knockout lines were grown in 15 cm culture plates to confluency. Cells were rinsed twice with 5 mL PBS at room temperature and incubated with 3 mL of 4 mM unloaded methyl-α-cyclodextrin (699020-02-5, Bio-Reagent) in PBS for 8 min at room temperature. The supernatant was collected, avoiding detached cells during collection. The residual cells were scraped into 2 mL PBS. Both fractions were frozen and stored at -80 °C until lipid extraction.

Aliquots of 200 µL from each fraction were vacuum-dried. Lipids were extracted with 695 µL methyl tert-butyl ether (MTBE)/methanol (10:3, v/v) and 5 µL SPLASH Lipidomix (Avanti Polar Lipids, Alabaster, AL), with mixing at 850 rpm for 1 h at 4 °C. After addition of 140 µL $H_2O$, samples were incubated for 15 min at 4 °C and centrifuged at 13,400 rpm for 15 min at 4 °C. The upper organic phase was collected and evaporated. Extracts were reconstituted in 200 µL isopropanol/methanol/chloroform (4:2:1, v/v/v) containing 7.5 mM ammonium formate.

Mass spectrometry was performed on a Q Exactive instrument (Thermo Fisher Scientific, Bremen, Germany) equipped with a TriVersa NanoMate robotic nanoflow ion source (Advion BioSciences, Ithaca, NY) and nanoelectrospray chips with a nozzle diameter of 4.1 µm. The ion source was controlled using Chipsoft 8.3.1. The ionization voltage was +0.96 kV in positive mode and -0.96 kV in negative mode, with a backpressure of 1.25 psi in both modes. The ion transfer capillary temperature was 200 °C and the S-lens RF level was 50%. Each acquisition lasted 14 min.

Full FT-MS spectra were acquired over m/z 300-1000 in positive mode from 0 to 0.2 min and negative mode from 7.7 to 7.9 min. The resolution was 140,000 at m/z 200, the automatic gain control (AGC) target was $3 \times 10^6$ and the maximum injection time was 3000 ms. Targeted selected ion monitoring was performed in positive mode from 1.7 to 7.5 min and negative mode from 7.9 to 14 min. The resolution was 140,000 at m/z 200, with an AGC target of $5 \times 10^4$, a maximum injection time of 650 ms and an isolation window of 20 Th. Target masses ranged from m/z 305 to 1005 in 10 Th increments. Free cholesterol was quantified by parallel reaction monitoring FT-MS/MS in positive mode from 0.2 to 1.7 min. The precursor isolation window was 0.8 Da, the normalized collision energy was 12.5%, the AGC target was $5 \times 10^4$ and the maximum injection time was 3000 ms, with one microscan.

Spectra were preprocessed using the repetition-rate filtering software PeakStrainer and stitched with an in-house script. Lipids were identified using LipidXplorer and molecular fragmentation query language queries for PC, PC O-, LPC, LPC O-, PE, PE O-, LPE, PI, PG, PA, PS, SM, TG, DG, Cer, Chol and CE. Intact masses were matched within a mass tolerance of ≤5 ppm. Lipid amounts were quantified by comparing isotopically corrected molecular ion abundances with those of internal standards of the same lipid class. An in-house R script excluded lipids detected only in blanks and required abundances above five times the blank and detection in at least 50% of samples. Supernatants and residual cells were filtered separately for the primary analysis. A dataset filtered jointly across both fractions was used for an additional composition comparison.

Data were analyzed in Python. Technical injections were averaged per sample, species amounts were summed per lipid class, and amounts measured in 200 µL aliquots were scaled to the fraction volumes of 3 mL (supernatant, cyclodextrin-extracted lipids) and 2 mL (residual cells) for comparisons between fractions. To estimate serum carry-over, the FBS used in the culture medium was analyzed by shotgun lipidomics at 2 µL and 20 µL. CE species detected in FBS and cells, making up at least 1% of CE in either and at least four-fold more abundant in FBS than in cells, were used as serum signature without reference to the extracts, and the volume of culture medium per dish that would supply all CE of an extract was calculated from the CE content of FBS and the 10% FBS in the medium.

Immunofluorescence Staining of GLUT1

Immunofluorescence stainings were performed in 96-well plates (655891, Greiner Bio-One) coated with fibronectin (20 µg/ml for 45 min at 37°C) (F4759, Sigma-Aldrich). Cells were seeded one day prior to staining with 15000 cells/well. Cells were washed three times with PBS and subsequently fixed with 4 % paraformaldehyde (30525-89-4, TCI)(in PBS) for 15 min at RT. Fixative was removed by washing three times with PBS and cells were permeabilized with 0.5 % Triton X-100 (37240.0, Serva)(in PBS) for 15 min at RT. The triton solution was removed and afterwards cells were incubated with blocking solution (0.1 % Triton, 2 % BSA (A8806, Sigma)) for 1 h at RT. The GLUT1 antibody (ab115730, abcam) was diluted 1:200 in blocking solution and incubated over-night at 4°C.

Cells were washed three times with PBS and afterwards treated with secondary antibodies and conjugated compounds (in blocking solution) for 1 h at RT. Cells were washed three times with PBS and counter-stained with 300 nM DAPI solution (D9542, Sigma-Aldrich)(in $H_2O$) for 5 min at RT. DAPI solution was removed by washing three times and cells were kept in PBS for imaging. Z-stacks were acquired on a single photon point scanning confocal system (Zeiss LSM 880 Airy inverted) using Zeiss Plan-Apochromat 63x 1.4 Oil DIC objective. An in-house Fiji[18] macro (consists of 3 individual parts) was used to analyze the images.

Imaging of Bifunctional Lipid Probes

Three bifunctional lipid probes were imaged independently: the long chain bifunctional phosphatidylethanolamine PE(18:1/$Y_{16}$), the short chain bifunctional phosphatidylethanolamine PE(14:0/$Y_{14}$), and a bifunctional phosphatidylcholine probe, PC(18:1/$Y_{16}$), described previously[17]. Liposomes were prepared by resuspending a final concentration of 0.75 mM of the respective bifunctional lipid, 0.375 mM Cholesterol (57-88-5, Sigma-Aldrich) and 0.375 mM POPC (26853-31-6, Avanti) in PBS. The suspension was extruded with 0.1 µm PC membranes (610005, Avanti). Liposomes were stored at 4°C for maximum 1 week before usage.

HCT116 WT cells and the ATP11C, ATP8B1 and TMEM30A knockout lines were seeded two days prior to the experiment with 5000 cells/well into 96-well plates (655891, Greiner). Liposomes were diluted to a final total lipid concentration of 0.5 mM in FBS free McCoy medium supplemented with 4 mM alpha-methyl Cyclodextrin. This mixture was incubated at 37°C for 30 min prior to incubation with the cells. Cells were washed with FBS free McCoy medium prior to adding the liposome mixture. Cells were incubated with the liposome mixture for 3 min at 37°C and subsequently washed with McCoy medium. After an additional chase time of 0, 1, 2, 7 or 27 min, samples were exposed to 10s of 365 nm UV light from an LED photoreactor (VS5252C45L9-

365, Violumas) and immediately fixed using 4 % PFA (30525-89-4, TCI) and 0.1 % glutaraldehyde (111-30-8, AppliChem) in PBS for 20 min at RT. Cells were washed with 100 mM Glycine solution in PBS and permeabilized using 0.1% Triton in PBS for 30min at RT. Each condition was acquired in two wells with several fields per well.
For immunofluorescence, the samples were blocked for 15min at RT using a 2 % BSA (9048-46-8, Sigma) in PBS solution. The plasma membrane was labelled in live cells before fixation using the MemBrite Fix 640/660 Cell Surface Staining Kit (30097, Biotium) as described by the manufacturer. The ER was labelled with a mix of rabbit antibodies against BAP31 (ab109304, abcam), calreticulin (ab2907, abcam), RTN4 (10950-1-AP, Proteintech) and lamin B1 (ab16048, abcam), each diluted 1:500. Endolysosomes were labelled with a mix of mouse antibodies against Rab5 (610725, BD Transduction Laboratories; 1:300), EEA1 (610457, BD Transduction Laboratories; 1:250), Rab7 (95746S, Cell Signaling Technology; 1:1000) and LAMP1 (sc-20011, Santa Cruz Biotechnology; 1:750). Primary antibodies were detected with Alexa Fluor-conjugated goat anti-rabbit and goat anti-mouse secondary antibodies (Thermo Fisher Scientific), diluted 1:2000.
The samples were washed with a 100 mM HEPES solution. The samples were treated with a click solution composed of 2 µM Azide dye (CLK-1296, Jena Bioscience), 5 mM L-ascorbic acid (95210, Merck), 0.5 mM THPTA (CLK-1010, Jena Bioscience) and 0.1 mM $CuSO_4$ (7758-98-7, ACROS OrganicsTM) in 100 mM HEPES for 30 min at 37°C. The click reaction was repeated one time with briefly washing with 100mM HEPES in between steps. Subsequently, the samples were incubated for 15 min at RT with a 1:5000 Hoechst (62249, Thermo Scientific) solution (in PBS).
Z-stacks were acquired with constant settings on an Olympus IX83 with a Yokogawa W1 CSU with a SoRa Disc using an Olympus 150X 1.45 NA Oil objective. Four channels were recorded at 0.4 µm z-steps: plasma membrane marker (640 nm excitation), lipid probe (561 nm), ER marker (488 nm) and endolysosome marker (405 nm).

Quantification of Organelle Partitioning

Image analysis was performed in Python using an in-house script. After camera offset subtraction, the plasma membrane, ER and endolysosome marker channels were denoised and segmented: the plasma membrane with an intensity threshold relative to each image combined with ridge detection, the ER by Otsu thresholding and endolysosomes by top-hat filtering and watershed separation of puncta. The marker channels were registered to the lipid channel, which was not denoised. Within the cell body region, defined by the lipid signal and the organelle masks, each pixel was assigned to a single compartment in the order endolysosome, plasma membrane and ER, and the remaining pixels to the cytoplasm. Lipid intensities were summed per compartment over all z-planes with signal, and the PM/(PM+ER) ratio was calculated per image. Individual images were analyzed (1492 images in total). Images identified as outliers by all four of the following tests were removed: modified z-score, Tukey fence, Grubbs test and generalized extreme studentized deviate test.

Transferrin Uptake Assay

Cells were seeded one day prior to staining with 4000 cells/well into 96-well plates (655891, Greiner). Transferrin uptake was determined by adding 20 µg/ml transferrin human serum, Alexa Fluor$^{TM}$ 647 conjugate (T23366, Invitrogen) to separate wells for 4 or 30 min of continuous uptake and subsequently cells were washed with PBS and fixed for 20 min with 4% PFA (30525-89-4, TCI). Fixed cells were counterstained with the ER and endolysosome antibody mixes described above for the imaging of bifunctional lipid probes. Z-stacks were acquired with constant settings on an Olympus IX83 with a Yokogawa W1 CSU with a SoRa Disc using an Olympus 150X 1.5 Oil objective. Transferrin puncta amounts and volume were analyzed with a python analysis script (see repository).

Cells were segmented with Cellpose-SAM in a composite of the ER and endolysosome channels at the z-plane of the nuclear equator, and cells touching the field border were excluded. The masks were applied to all z-planes. Transferrin structures were detected by thresholding a smoothed copy of the stack at the stack median plus three times the scaled median absolute deviation and separated by watershed segmentation; intensities were measured in the raw data. Structures of 5 to 500 voxels were counted as puncta and larger ones as large puncta, and the volume of the largest structure per cell is reported as the endocytic recycling compartment volume. Total transferrin per cell was measured as the offset-subtracted intensity integrated over the cell mask and all z-planes.

Individual cells from two biological replicates were analyzed (4851 cells from 280 fields). Each KO line was compared with WT at the same uptake time by a two-sided Mann-Whitney U test with Benjamini-Hochberg correction.

Assessment of Lipid Droplet Morphology

Oleic acid (O1383, Merck) was warmed to 37°C until it was completely liquified. Subsequently, the oleic acid was diluted to 150 mM in 50 % EtOH. A 100 mg/mL solution of fatty acid free BSA (A8806, Merck) was prepared in $H_2O$ and mixed by pipetting with equal volumes with the oleic acid dilution. The mixture was incubated for 1h at 37°C. The resulting cloudy and viscous oleic acid/BSA complex was mixed 1:300 in complete cell culture medium to a final concentration of 0.25 µM oleic acid.

Cells were seeded one day prior to staining with 20000 cells/well into 96-well plates (655891, Greiner). Cells were incubated with the prepared oleic acid/BSA mixture O/N. Afterwards, cells were washed three times with PBS and medium was added and left on the cells for 8 h. The experiment was performed in three biological replicates with the following conditions: untreated, O/N treated and 8 h after treatment. Cells were fixed with 4 % PFA and stained with a 1:5000 Hoechst (62249, Thermo Scientific) solution (in PBS). Afterwards, LipidSpot610 (70069, Biotium) was added to the cells without subsequent washing steps. Z-stacks were acquired on a single photon point scanning confocal system (Zeiss LSM 880 Airy inverted) using Zeiss Plan-Apochromat 63x 1.4 Oil DIC objective. Lipid droplet volumes and numbers were analyzed with an in-house Fiji[18] macro (consists of 3 individual parts, can be found on the repository).

Seahorse Assays

One day prior to experiments, 5000 cells were seeded into XFe96/XF cell culture microplates. XFe/XF96 analyzer was used to obtain oxygen consumption rate and extracellular acidification rates prior and after addition of inhibitors. Experiments were carried out following vendor's protocol. For the experiments the following kits were used: Seahorse XF Glycolytic Rate Assay Kit (Agilent, 103344-100) and Seahorse XF Real-Time ATP Rate Assay Kit (Agilent, 103592-100). After the end of experiments, cells were fixed with 4% PFA for 15 minutes, then washed with PBS and stained with DAPI 1ug/ml and stored at 4C until further processing. Images were acquired using the Yokogawa CV7000 automated microscope platform for 9 fields per well for 1 channel (DAPI) using a 10x objective (NA= 0.75). Nuclei were segmented from these images using a CellProfiler[19] pipeline and data was further processed with KNIME (version 4.6) to calculate the effective number of cells per well, which was then used to normalize the measured metabolic rates.

Assessment of ATP Depletion Dynamics

One day prior to experiment, 5000 cells (WT and respective KOs) were seeded into 18-well dishes (81817, ibidi). After 6 h, cells were transfected with cyto-iATPSnFR1.0[20] plasmid using X-tremeGENE™ HP DNA Transfection Reagent (6366244001, Merck/Roche) with a DNA (weight):reagent (volume) ratio of 1:3. Right before imaging, 50 µl of imaging buffer (20 mM HEPES, 115 mM NaCl, 1.2 mM $CaCl_2$, 1.2 mM $MgCl_2$, 1.2 mM $K_2HPO_4$, 20 mM glucose) was added to the cells. Fluorescence intensity of the ATP sensors was imaged over time and after 96 sec 50 µl of IB without glucose containing 20 mM 2-Deoxy-D-glucose (D8375, Merck), 20 µM Antimycin A (A8674, Merck) resulting in final concentrations of 10 mM 2-DG and 10 µM Antimycin A. To provide blank baseline data for the addition experiments, identical volume of the imaging buffer was added to the cells. Fluorescence intensity was imaged every 5 sec for 10 min. Images were acquired on a laser scanning confocal (Olympus FluoView3000) using a UPLXAPO40x0 40x oil (NA = 1.4) objective. Image analysis was performed in python and the script is available on the repository. In brief: Cells were segmented using mean thresholding (skimage.filters.threshold_mean) followed by watershed segmentation (touching_objects_labelling as implemented in napari_simpleitk). Fluorescence intensities were extracted for individual cells and bleach corrected using data from untreated conditions. Bleach corrected data were normalized to initial fluorescence (ΔF/F0). For each cell, the initial velocity of fluorescence decrease was estimated by fitting a linear regression to time points from compound addition (96s) until one-third of the time to reach the fluorescence minimum. This captures the early phase of the response before nonlinear effects dominate.

Nucleotide Measurements

Cells were seeded two days prior to proteomics sample collection with 5 x $10^5$ cells in 60 mm Nunclon™ Delta Surface dishes (150326, Thermo Scientific). Cells were washed twice with ice-cold PBS and subsequently scraped (3010, Corning) in 300 µL MeOH/H2O (80:20, v/v) and immediately snap-frozen and stored at -80°C until processing. For internal standards: Adenosin-$^{13}C_{10}$ 5′-triphosphat (741167, Sigma), Adenosine-15N5 5′-diphosphate (741167, Sigma), Adenosin-13C10,15N5-5′-monophosphat (650676, Sigma) were used. Subsequently, the mixture subjected to three cycles of snap-freezing and thawing and afterwards centrifuged at 13000 x g for 10 min. The supernatant was transferred to a glass vial and the LC-MS/MS analysis was performed using liquid chromatography-tandem mass spectrometry on an ultra-performance liquid chromatography system (Aquity I-class, Waters) coupled to a triple quadrupole linear ion trap mass spectrometer (QTRAP 5500, Sciex).

For normal phase chromatography, an XBridge BEH Amide Column, 130Å, 3.5 µm, 2.1 mm × 100 mm from Waters was used. The mobile phase consisted of eluent A (95% acetonitrile, 10 mM ammonium acetate, and 0.01% $NH_4OH$) and eluent B (40% acetonitrile, 10 mM ammonium acetate, and 0.01% $NH_4OH$). Chromatographic separation was achieved at 40°C with the

following gradient program: Eluent B, from 0% to 100% within 18 minutes; 100% from 18 to 21 minutes; 0% from 21 to 26 minutes. The flow rate was set at 0.300 mL/min.
The metabolites were analyzed in multiple reaction monitoring (MRM) scan mode using negative electrospray ionization (ESI). The ion source parameters were as follows: curtain gas (40 psi), ESI voltage (-5500V), source temperature (500°C), gas 1 (70 psi), and gas 2 (50 psi). Compound-dependent source and fragmentation parameters were set to a 100V declustering potential, 10V entrance potential, 45V collision energy, and 10V cell exit potential.
Data acquisition was performed using Analyst 1.7 (Sciex), and data processing was done using the Sciex OS-MQ software package. Internal standards, and protein amounts (BCA) were used for quantification.

Calorimetry Experiments presented in Fig. 5E-H
For these experiments, cells were grown in McCoy's supplemented with glutamine instead of GlutaMAX without the addition of antibiotics. Calorimetry measurements were done using TAM IV isothermal microcalorimeter (TA Instruments) in either "mini" module (with 6 channels with built-in reference) or "nano" module (paired sample and reference channel). Two controls and 4 samples were always used in the "mini" module. 4-mL glass vials with 1 $cm^2$ horizontal growth surface were coated for 1 h at 37 °C with 180 µL of 10 µg/mL fibronectin in PBS (5050, Advanced Biomatrix) and washed once with PBS. Cells were seeded from premixed suspension of desired concentration to reach density of 25,000 cells/$cm^2$ and total volume of 2 mL. Control vials contained medium only. All vials were closed with aluminum caps with crimping tool using maximal force and hooks for sample holder rods were screwed into the caps.
Experiments were carried out according to the experimental wizard in the acquisition software TAM Assistant (TA Instruments, version v2.0.175.1). The thermostat of the calorimeter oil bath was set to 37.0 °C. Once equilibrated, gain calibration was performed for all modules. Upon starting an experiment, baseline of heat flow was captured for 30 min and subtracted as an offset. The samples were then inserted halfway into equilibration position for at least 15 min, before lowering all the way down to measuring position. Signal was considered correct 45 min after. Heat data were collected until cells reached saturation, at least for 3 days. Samples were then removed and final baseline collected for 1 hour after all channels equilibrated again.

Calorimetry Experiments presented in Fig. 5I
HCT116 cells were trypsinised and 100000 cells were seeded into a 4 ml glass vial (Waters) in a total volume of 2 ml cell culture medium. Prior to seeding, glass vials were coated with fibronectin (20 µg/ml for 45 min at 37°C). Additionally, to determine protein concentration at different time points, two vials for every time point were prepared. Vials were closed with lids (Waters) and placed for 30 min in the incubator. Determined vials for measuring were placed in the "nano" module of TAM IV calorimeter (Waters/TA Instruments) and the heat flow and cumulative heat were measured over the course of 16 h. Vials for protein concentration determination were left (with closed lid) in the incubator and two random vials were picked every four hours for determination of protein concentration. Cells were washed three times with ice cold PBS and 1 ml of RIPA buffer (89901, Thermo Scientific) was added for 5 min. Afterwards, cells were scraped and transferred to microcentrifugation tubes and centrifuged at 14000 x g for 15 min. The supernatant was transferred to a new tube and protein concentration was

determined by BCA (23235, Thermo Scientific). For the calculations of total protein concentration, the protein concentration determined in the reference vial (just medium) was subtracted.
Experiments were carried out according to the experimental wizard in the acquisition software TAM Assistant (TA Instruments, version v2.0.175.1). The thermostat of the calorimeter oil bath was set to 37.0 °C. Once equilibrated, gain calibration was performed for all modules. Upon starting an experiment, baseline of heat flow was captured for 30 min and subtracted as an offset. The samples were then inserted halfway into equilibration position for at least 15 min, before lowering all the way down to measuring position. Signal was considered correct 45 min after. Afterwards, data was plotted in R.

<u>QPI experiments</u>
QPI experiments were done using Q-Phase quantitative phase microscope (Telight). Phase-contrast µ-Slide Ph+ dishes with glass bottom (2-well (80297) or 4-well (80447) dishes, Ibidi) were coated the same way as calorimetry vials (seen in Fig. 5E-H), using 1.5 mL of fibronectin solution. Cells were seeded in similar fashion as in calorimetry experiments; 1.5 mL of cell suspension with 25,000 cells/cm$^2$ final density were seeded with SPY555 DNA dye added to final concentration 1× for enhancing cell segmentation (SC201, Spirochrome). Samples were sealed with 24×60 mm coverslips with dentistry-grade silicone used as glue. Microscope was tempered to 35.3 °C inside a temperature-controlled box; the sample itself was brought to 37 °C by adding extra transparent heater from top (also to prevent condensation inside the sealed sample).
All imaging was done with 10× objective. Prior to each experiment, fresh background was captured. Sample was then inserted onto stage and secured in custom-made aluminum frame with magnets to prevent drift. Tuning of QPI signal was first done using automated wizard within the SophiQ acquisition program (Telight, version 10.1.5.gb1cbb). Each field of view (FOV) was fine-tuned manually for maximal signal value. QPI holograms and fluorescence with TRITC filter were acquired with 8 min acquisition interval over ≥48 h with ≥5 FOVs per well. Signal values were checked throughout the experiment; when necessary, imaging was stopped and all FOVs retuned.
*QPI data processing*
Phase images were calculated post-hoc from captured raw QPI holograms. Phase and fluorescence images were exported as .tiff with original units (pg/µm$^2$ and arbitrary gray values, respectively). Fluorescence images were cropped and shifted to match and overlay phase images (1200×1200 pixels ≌ 592×592 µm$^2$ FOV). Segmentation of images was done using Cellpose[21] on phase channel with nuclei in fluorescent channel as auxiliary cue; cells touching edges were removed. Custom trained model for segmentation was used. Properties of segmented cells were analysed and collected into .csv table using Python3 and libraries pandas, numpy, scikit-image, and visualised using plotly.
Average cell mass was calculated for each FOV and subsequently average total cell mass of all FOVs in a sample was calculated. Mass data were recalculated to µg/cm$^2$ based on the size of the FOV.
*Mass-specific heat dissipation plots*
Mass-specific heat dissipation plot (Fig. 5H) shows complete averaged QPI data and complete averaged calorimetry data for each cell line. Data were cropped to 250 mJ and 175 µg to show

the initial linear regime, which was fitted with least squares polynomial fit of degree 1. The errors were calculated as the standard deviation of the distribution of slope fits from bootstrapped distribution performed on 20 % of randomly selected points (changed with each resample) with 10,000 resamples.

Data availability, presentation and processing statement

All raw data and images can be found on the following repository: EDMOND (https://doi.org/10.17617/3.1HISIJ).

Lipidomics data presented in Fig. 2 and Extended Data Fig. 3 was acquired in 2 batches, each containing a WT sample for comparison. In all analyses, the respective WT of the batch was compared to the KO cells except for Fig. 2A, where a general overview for both batches is shown as an average.

For fluorescence microscopy, laser settings were always the same in the respective experiments between the individual KO lines. Only for representation purposes, contrast and brightness was adjusted in the same manner within the respective experiments.

For lipid droplet and GLUT1 analyses Shapiro-Wilk-tests were performed to determine if data were normally distributed. In this was not the case, a two-tailed Mann-Whitney-U test with a significance level alpha 0.01 was used to test $H_0$ hypotheses.

Chemical Synthesis

All chemicals were obtained from commercial sources (Acros, Sigma-Aldrich, TCI chemicals, Alfa Aesar, Roth, Fluka or Merck) and were used without further purification. Solvents for flash chromatography were obtained from VWR and dry solvents were obtained from Sigma. Deuterated solvents were obtained from Deutero GmbH, Karlsruhe, Germany. TLC was performed on precoated plates of silica gel (Merck, 60 F254) using UV light (254 or 365 nm) or a solution of phosphomolybdic acid in EtOH (10 g phosphomolybdic acid, in 100 mL EtOH) for analysis. Preparative column chromatography was performed using silica gel from Merck, Darmstadt, Germany (silica 60, grain size 0.063-0.200 mm) with a pressure of 1 bar. Detailed purification conditions are given for the respective compounds. $^{1}$H,$^{13}$C and $^{31}$P NMR-spectra were measured on 400 MHz AdvanceTM III HD Nanobay Bruker spectrometer. Chemical shifts of $^{1}$H- and $^{13}$C-NMR-spectra are referenced indirectly to tetramethylsilane. J values are given in Hz and chemical shifts in ppm. Splitting patterns are designated as follows: s, singlet; d, doublet; t, triplet; q, quartet; m, multiplet; $m_c$, centered multiplett. $^{13}$C-NMR-spectra were broadband hydrogen decoupled. Mass spectra were recorded using a QExactive Orbitrap FTMS instrument (ThermoFisher Scientific) equipped with a robotic nanoflow electrospray ion source. FTMS were acquired for the mass range of m/z 200-1500 with the target mass resolution set to 140000 at m/z 200. Samples were dissolved in a solution of iPrOH/MeOH/CHCl3 4:2:1 containing 7.5 mM ammonium formiate. The spectra were evaluated using the Xcalibur Qual Browser software.

(2R)-3-(((2-((((9H-fluoren-yl)methoxy)carbonyl)amino)ethoxy)(hydroxy)phosphoryl) oxy)-2-hydroxypropyl oleate (**P1**)

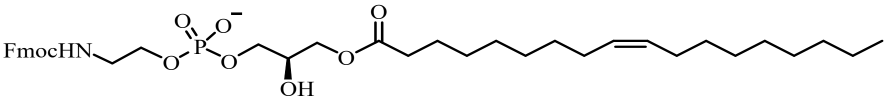

**P1**

200.0 mg of Lyso-PE 18:1 (417.0 µmol, 1 eq.) were dissolved in 6 ml dry chloroform and activated with 106.6 µl of DIPEA (79.1 mg, 612.0 µmol, 1.5 eq.) for 15 minutes. Subsequently, a solution of 115.9 mg FmocCl (448.0 µmol, 1.1 eq.) in 3 ml dry chloroform was added and the reaction mixture was stirred for 22 h under inert gas atmosphere. After removal of the solvent the crude product was purified by flash chromatography (eluent chloroform/ MeOH/ water 65/35/2). Fmoc-Lyso-PE 18:1 (**P1**) was isolated as a white solid.

$^{1}$H NMR (400 MHz, MeOD) δ = 7.77 (d, $J$ = 7.5 Hz, 2H), 7.64 (d, $J$ = 7.4 Hz, 2H), 7.36 (dd, $J_1$ = 7.4 Hz, $J_2$ = 7.4 Hz ), 7.29 (dd, $J_1$ = 7.4 Hz, $J_2$ = 7.4 Hz ), 5.31 ($m_c$, 2H), 4.28 (d, $J$ = 14.6 Hz, 2H), 4.24 – 4.01 (m, 3H), 4.00 – 3.82 (m, 5H), 3.36 (t, $J$ = 5.2 Hz, 2H), 2.28 (t, $J$ = 7.5 Hz, 2H), 2.05 – 1.89 (m, 4H), 1.54 ($m_c$, 2H), 1.40 – 1.19 (m, 20H), 0.88 (t, $J$ = 6.8 Hz, 3H) ppm.
$^{13}$C NMR (101 MHz, MeOD) δ = 175.30, 158.85, 145.28, 142.55, 130.84, 130.77, 128.77, 128.15, 126.22, 120.92, 69.86, 67.92, 67.56, 66.20, 65.47, 42.64, 34.86, 33.04, 30.82, 30.78, 30.59, 30.43, 30.32, 30.30, 30.18, 28.11, 25.93, 23.72, 14.48 ppm.*
*[1]signals at 69.8, 67.5 65.4 and 42.6 ppm appear as two peaks because of the presence of two confomers
*[2]Signals between 30.2 and 30.8 ppm overlap, not all signals resolved individually
$^{31}$P NMR (162 MHz, MeOD) δ = -1.41 ppm.
HR-MS (ESI negative) m/z calculated for $C_{38}H_{56}NO_9P$: 701.36927; found: 700.363 $[M-H]^-$.

<u>(2R)-3-(((2-aminoethoxy)(hydroxy)phosphoryl)oxy)-2-((6-(3-(non-8-yn-1-yl)-3H-diazirin-3-yl)hexanoyl)oxy)propyl oleate (**P2, PE(18:1/$Y_{16}$)**)</u>

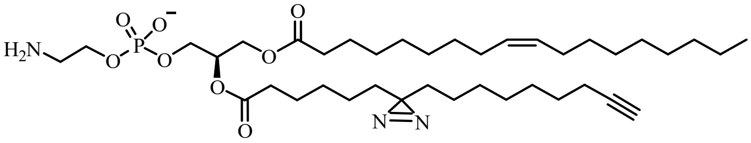

**P2**

57.0 mg of the bifunctional fatty acid (186 µmol, 1 eq.) were dissolved in 3 ml dry DMF and activated with 64.4 mg EDC·HCl (336 µmol, 1.8 eq.), 11.8 mg DMAP (96.6 µmol, 0.5 eq.) and 107 µl of DIPEA (614 µmol, 3.3 eq.) for 12 minutes. After adding a solution of 133 mg Fmoc-Lyso-PE 18:1 (**P1**) (190 µmol, 1 eq.) in 3 ml dry DMF and 5 ml dry chloroform, the reaction mixture was stirred for 19 h under an inert gas atmosphere. After removal of the solvent the product mixture was purified by flash chromatography (eluent chloroform/MeOH 70/30). Product-containing fractions were combined and used directly for the subsequent Fmoc-deprotection step.
The crude Fmoc-PE was dissolved in 3 ml dry DMF and 2 drops of piperidine were added. The reaction was stopped after 1.5 h by removal of the solvent. The crude product was purified by flash chromatography (eluent chloroform/ MeOH/ water 70/30/2). The bifunctional PE **P2** was isolated as yellowish oil.

$^{1}H$ NMR (400 MHz, MeOD) δ = 5.35 ($m_c$, 2H), 5.23 ($m_c$, 1H), 4.44 (dd, *J* = 12.0, 3.2 Hz, 1H), 4.18 (dd, *J* = 12.0, 6.7 Hz, 1H), 4.04 ($m_c$, 2H), 4.00 (t, *J* = 5.9 Hz, 2H), 3.17 (t, *J* = 4.9 Hz, 2H), 2.33 (t, *J* = 7.3 Hz, 2H), 2.32 (t, *J* = 7.4 Hz, 2H), 2.21 – 2.08 (m, 3H), 2.09 – 1.99 (m, 4H), 1.69 – 1.53 (m, 4H), 1.53 – 1.45 (m, 2H), 1.43 – 1.22 (m, 32H), 1.16 – 1.02 (m, 4H), 0.91 (t, *J* = 6.7 Hz, 3H) ppm. *[1]signals (triplets) at 2.33 ppm and 2.32 ppm overlap
$^{13}C$ NMR (101 MHz, MeOD) δ = 174.92, 174.45, 130.93, 130.79, 85.02, 71.92, 69.44, 64.93, 63.60, 62.95, 41.71, 41.65, 34.90, 33.83, 33.74, 33.07, 30.85, 30.84, 30.62, 30.46, 30.36, 30.33, 30.22, 30.19, 29.97, 29.69, 29.63, 29.50, 28.15, 26.00, 25.75, 24.87, 24.66, 23.75, 18.99, 14.48 ppm.*[2,3]
*[2]signals at 71.9, 64.9, 62.9 and 33.8 ppm appear as two peaks because of the presence of two confomers
*[3]Signals between 29.4 and 31.8 ppm overlap, not all signals resolved individually
$^{31}P$ NMR (162 MHz, MeOD) δ = 0.12 ppm.
HR-MS (ESI negative) m/z calculated for $C_{39}H_{70}N_3O_8P$: 739.49005; found: 738.482 $[M-H]^-$.

(2R)-3-(((2-((((9H-fluoren-9-yl)methoxy)carbonyl)amino)ethoxy)(hydroxy)phos-phoryl)oxy)-2-hydroxypropyl tetradecanoate (**P3**)

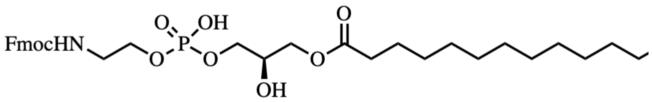
**P3**

170 mg Lyso PE 14:0 (400 µmol, 1 eq.) were dissolved in 6 ml dry chloroform and 3 ml dry DMF under an argon atmosphere. 100 µl DIPEA (600 µmol, 1.5 eq.) were added dropwise. After 15 min 114 mg FmocCl (440 µmol, 1.1 eq.) dissolved in 6 ml dry chloroform were added. The reaction mixture was stirred at room temperature overnight. The solvents were removed and the crude product was purified by flash chromatography (eluents chloroform/MeOH/water 65/35.1.5). Fmoc-Lyso-PE 14:0 (**P3**) was isolated light-yellow oil.

$^{1}H$ NMR (400 MHz, MeOD) δ = 7.79 (d, *J* = 7.5 Hz, 2H), 7.66 (d, *J* = 7.5 Hz, 2H), 7.38 (t, *J* = 7.4 Hz, 2H), 7.31 (t, *J* = 7.4 Hz, 2H), 4.32 (d, *J* = 7.1 Hz, 2H), 4.24 – 4.04 (m, 3H), 4.01 – 3.83 (m, 5H), 3.36 (t, *J* = 5.6 Hz, 2H), 2.29 (t, *J* = 7.5 Hz, 2H), 1.63 – 1.49 (m, 2H), 1.37 – 1.16 (m, 21H), 0.89 (t, *J* = 6.7 Hz, 3H) ppm.
$^{31}P$ NMR (162 MHz, MeOD) δ = 0.09 ppm.
$^{13}C$ NMR (101 MHz, MeOD) δ = 175.39, 158.88, 145.33, 142.58, 128.77, 128.17, 126.27, 120.91, 69.92, 69.85, 67.95, 67.55, 67.49, 66.23, 65.44, 42.72, 34.89, 33.08, 30.80, 30.77, 30.73, 30.61, 30.48, 30.42, 30.21, 25.96, 23.74, 14.45. ppm
HR-MS (ESI negative) m/z calculated for $C_{34}H_{50}NO_9P$: 647.3223; found: 646.313 $[M-H]^-$.

(2R)-3-(((2-aminoethoxy)(hydroxy)phosphoryl)oxy)-2-((6-(3-(hept-6-yn-1-yl)-3H-diazirin-3-yl)hexanoyl)oxy)propyl tetradecanoate (**P4, PE(14:0/$Y_{14}$)**)

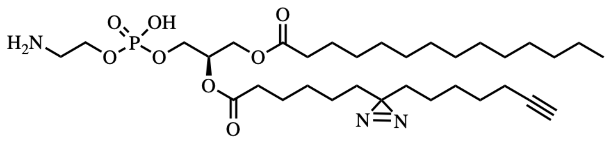

**P4**

86.5 mg of **P3** (133μmol, 1 eq.) were dissolved in 2 ml dry DMF and 2 ml $CDCl_3$. Bifunctional fatty acid was dissolved in 2ml $CDCl_3$ and activated with 66.5 mg EDC·HCl (347 μmol, 2.6 eq.) and 8 mg DMAP (66.5 μmol, 0.5 eq.) for 12 minutes. The activated fatty acid was added dropwise to **P3** and the mixture was stirred overnight at RT. The solvents were removed under reduced pressure and the product mixture was purified by flash chromatography (eluents 20% MeOH/chloroform). Product-containing fractions were combined and used directly for the subsequent Fmoc-deprotection step.

The solvent was removed and the crude product (63 mg) was dissolved in 3 ml dry DMF under an argon atmosphere. 2 drops of piperidine were added and the reaction mixture was stirred for 1.5 h at room temperature. The solvent was removed under reduced pressure and the crude product was purified by flash chromatography (eluents chloroform/MeOH/water 70/30/1). The bifunctional PE **P4** was isolated as yellowish oil.

$^{1}H$ NMR (400 MHz, MeOD) δ = 5.22 ($m_c$, 1H), 4.43 (dd, *J* = 12.1, 3.3 Hz, 1H), 4.18 (dd, *J* = 12.0, 6.6 Hz, 1H), 4.09 – 3.93 (m, 4H), 3.16 (t, *J* = 4.9 Hz, 2H), 2.33 (t, *J* = 7.4 Hz, 2H), 2.32 (t, *J* = 7.5 Hz, 2H), 2.21 – 2.10 (m, 3H), 1.66 – 1.52 (m, 4H), 1.52 – 1.42 (m, 2H), 1.42 – 1.22 (m, 30H), 1.18 – 1.04 (m, 4H), 0.90 (t, *J* = 6.6 Hz, 3H) ppm.
$^{13}C$ NMR (101 MHz, MeOD) δ = 174.90, 174.40, 84.81, 71.89, 71.81, 69.60, 64.92, 64.86, 63.56, 63.01, 62.95, 41.69, 41.63, 34.90, 34.88, 33.80, 33.68, 33.10, 30.84, 30.79, 30.66, 30.51, 30.47, 30.22, 29.68, 29.46, 29.36, 26.01, 25.75, 24.67, 24.65, 24.47, 23.76, 18.91, 14.49 ppm.
$^{31}P$ NMR (162 MHz, MeOD) δ = 0.12 ppm.
HR-MS (ESI negative) m/z calculated for $C_{33}H_{60}N_3O_8P$: 657.4118; found: 656.403 $[M-H]^-$.

<u>NMR Spectra of Synthesized Lipids</u>

Compound **P1** $^{1}$H spectrum

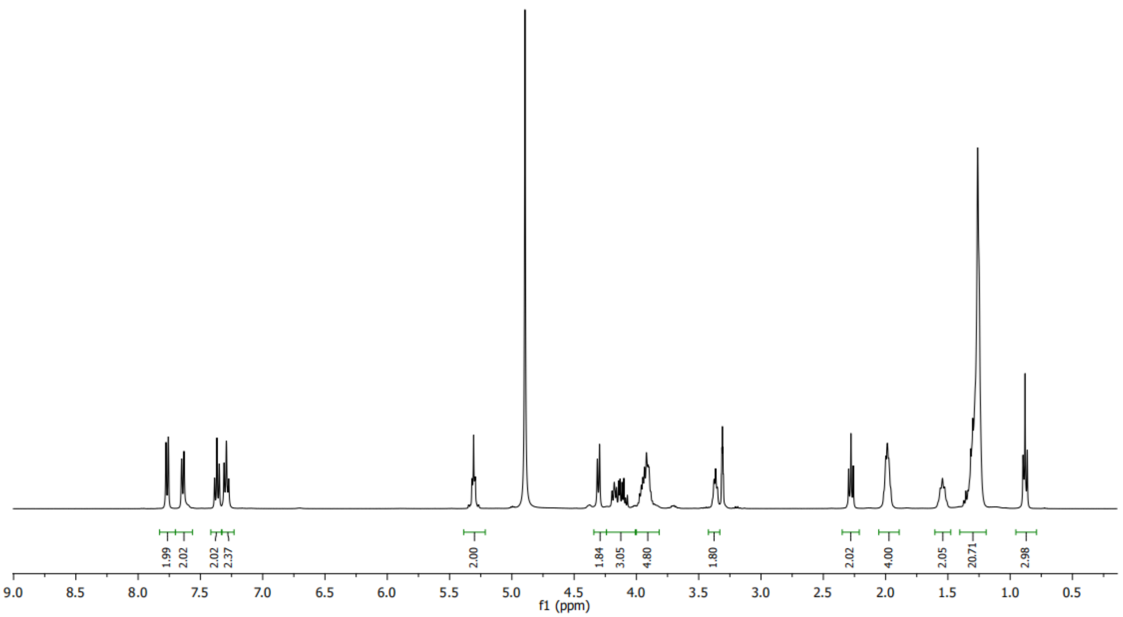


Compound **P1** $^{13}$C spectrum

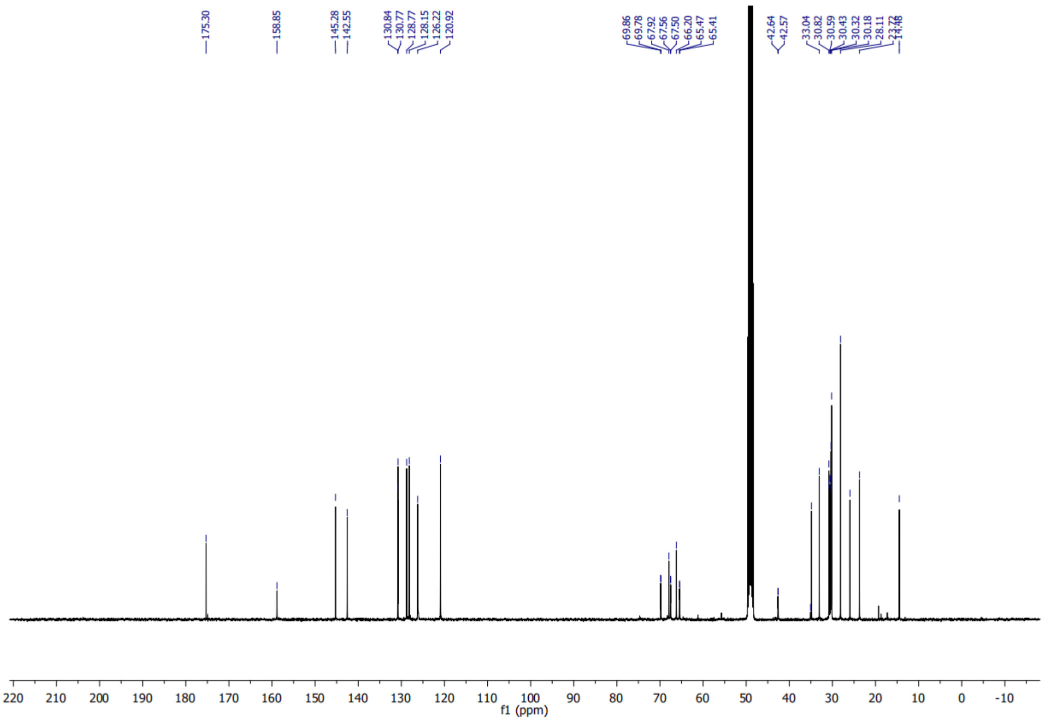

Compound **P1** $^{31}$P spectrum

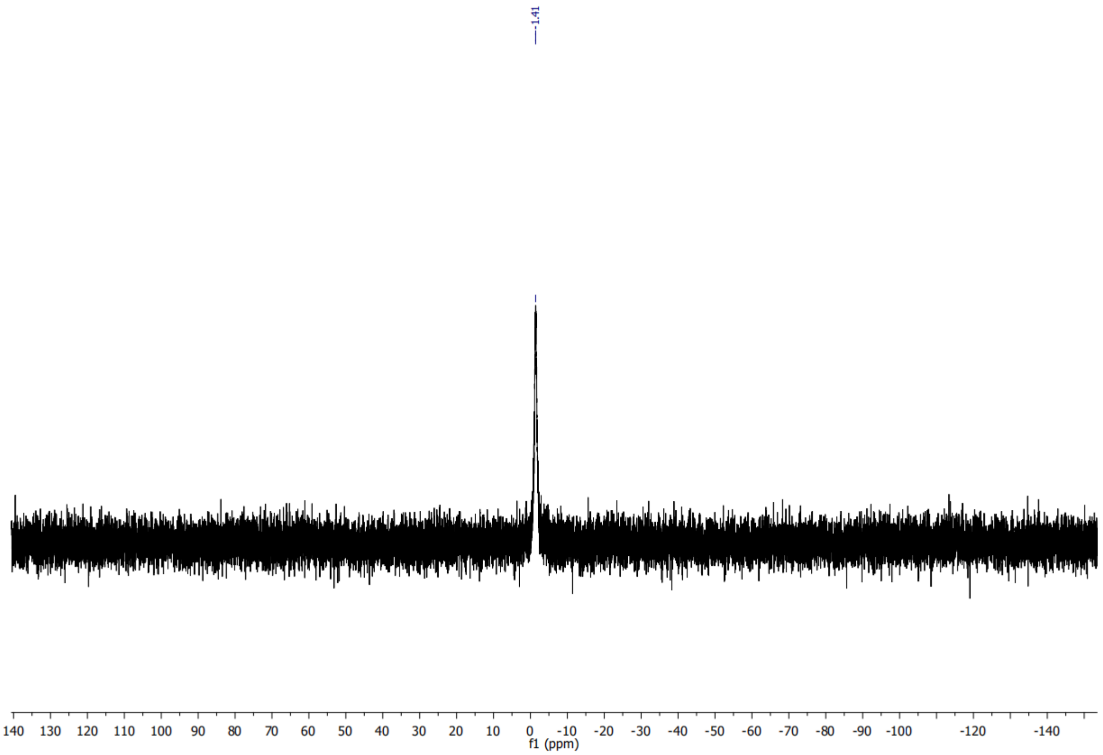


Compound **P2** $^{1}$H spectrum

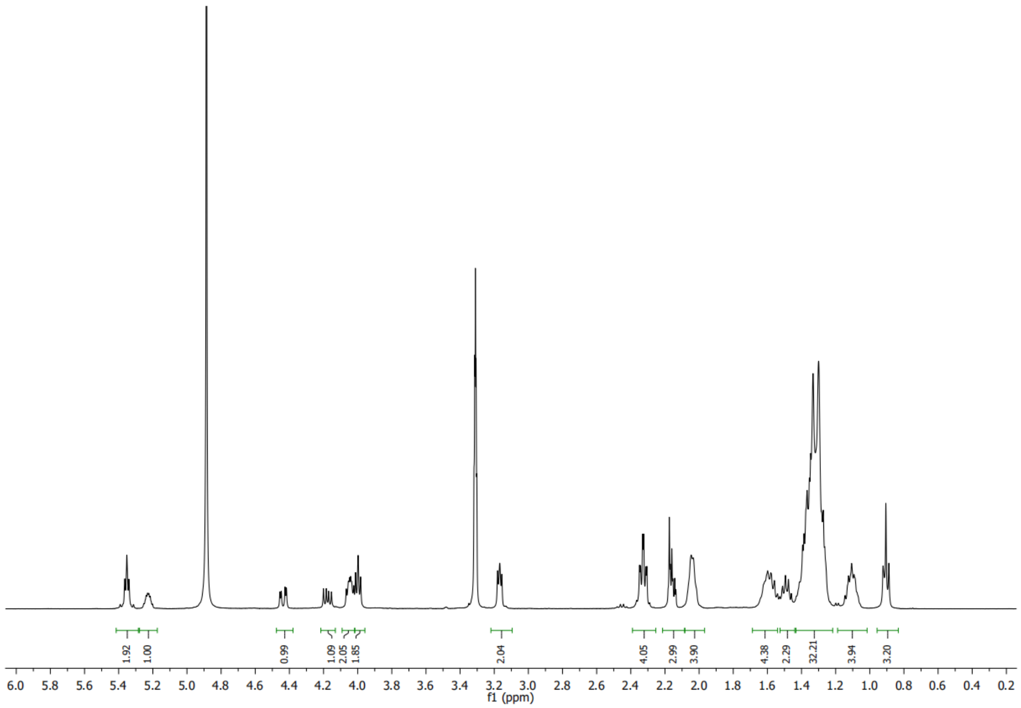

Compound **P2** $^{13}$C spectrum

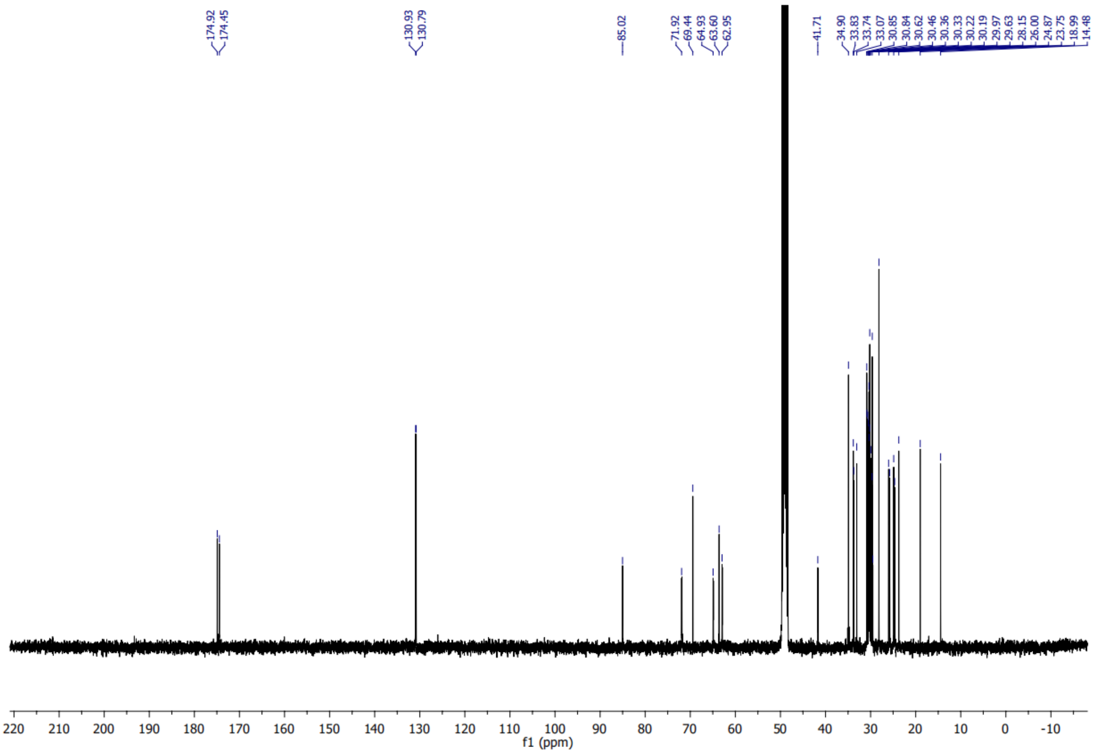


Compound **P2** $^{31}$P spectrum

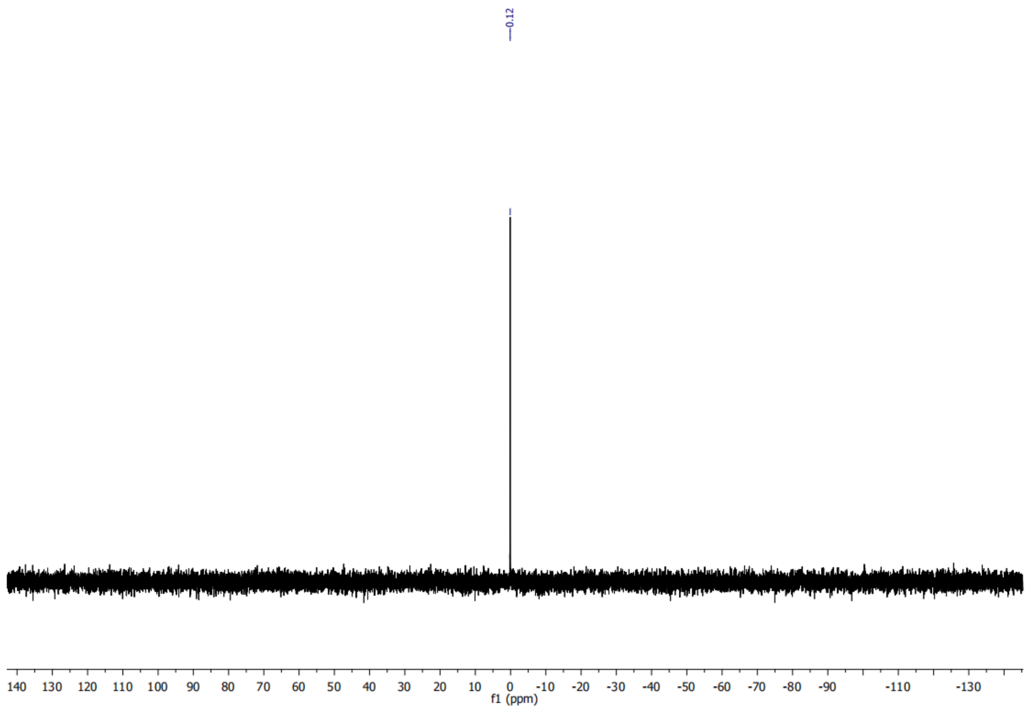

Compound **P3** $^{1}H$ spectrum

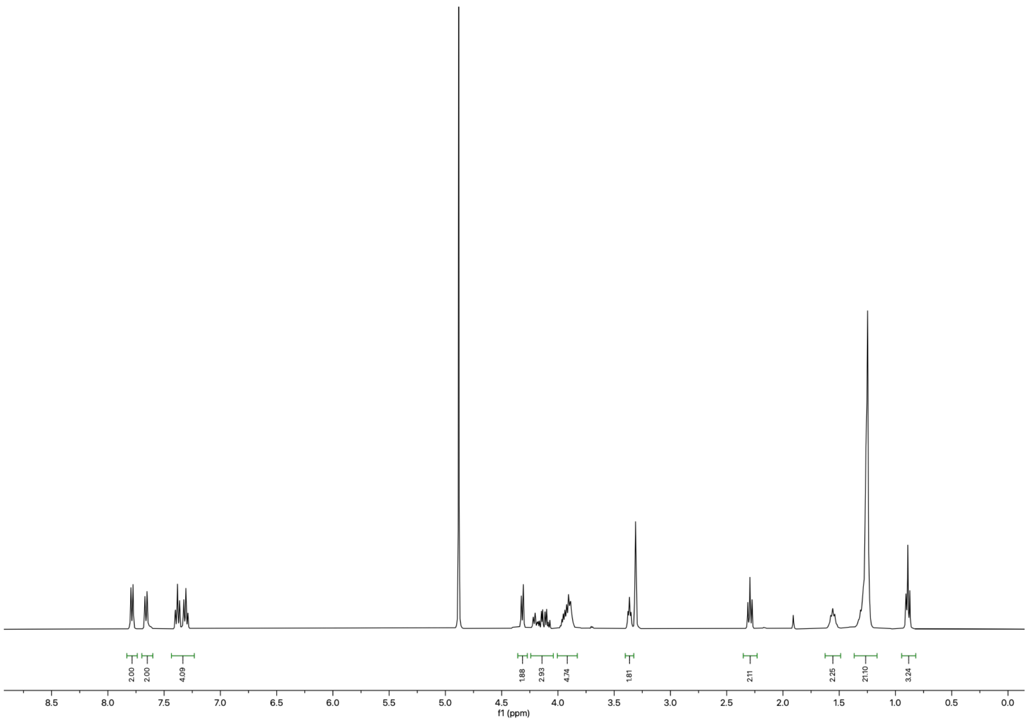

Compound **P3** $^{13}C$ spectrum

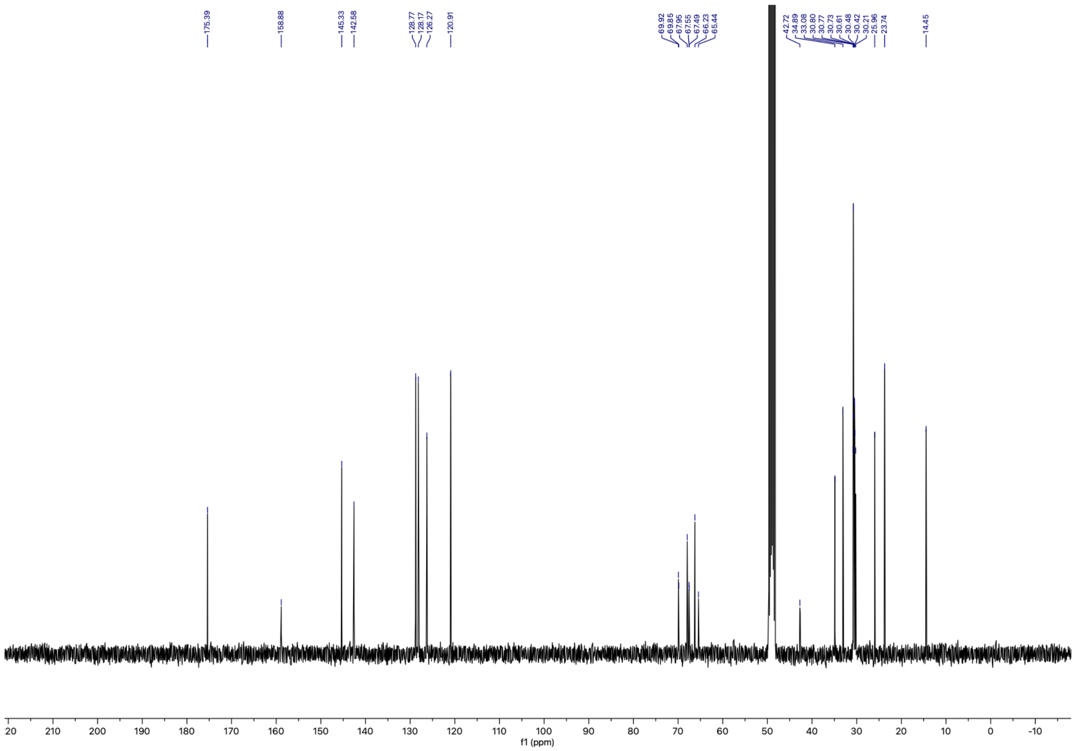

Compound **P3** $^{31}$P spectrum

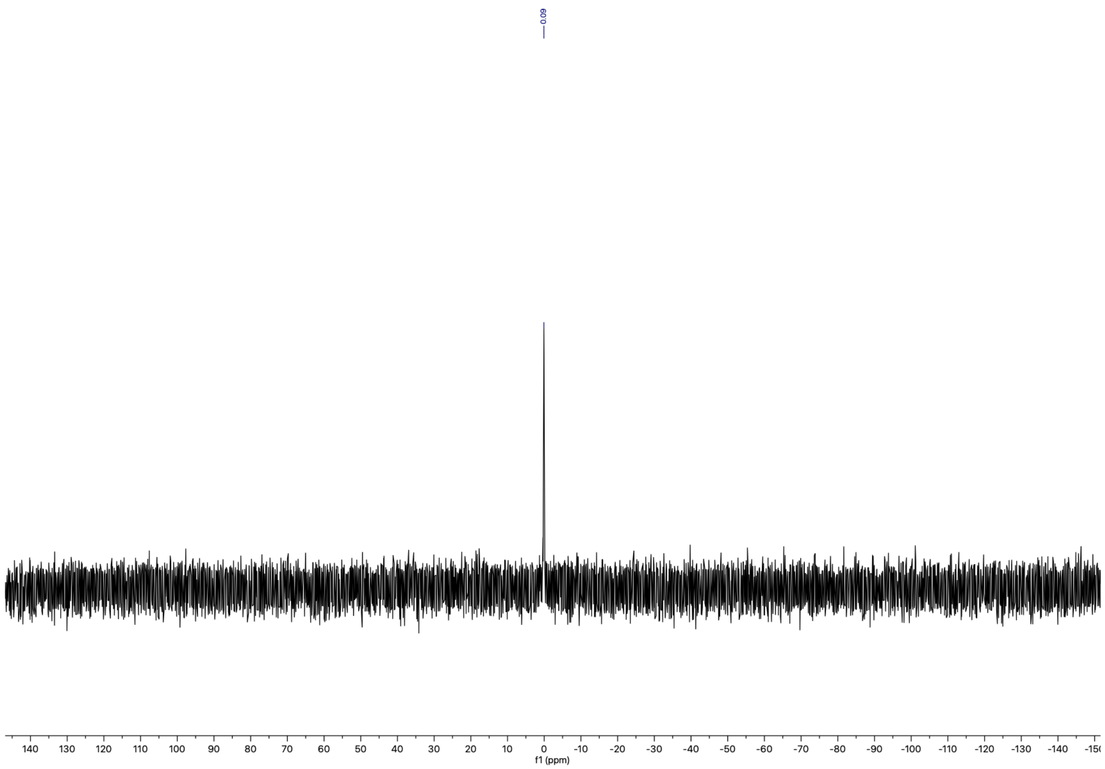


Compound **P4** $^{1}$H spectrum

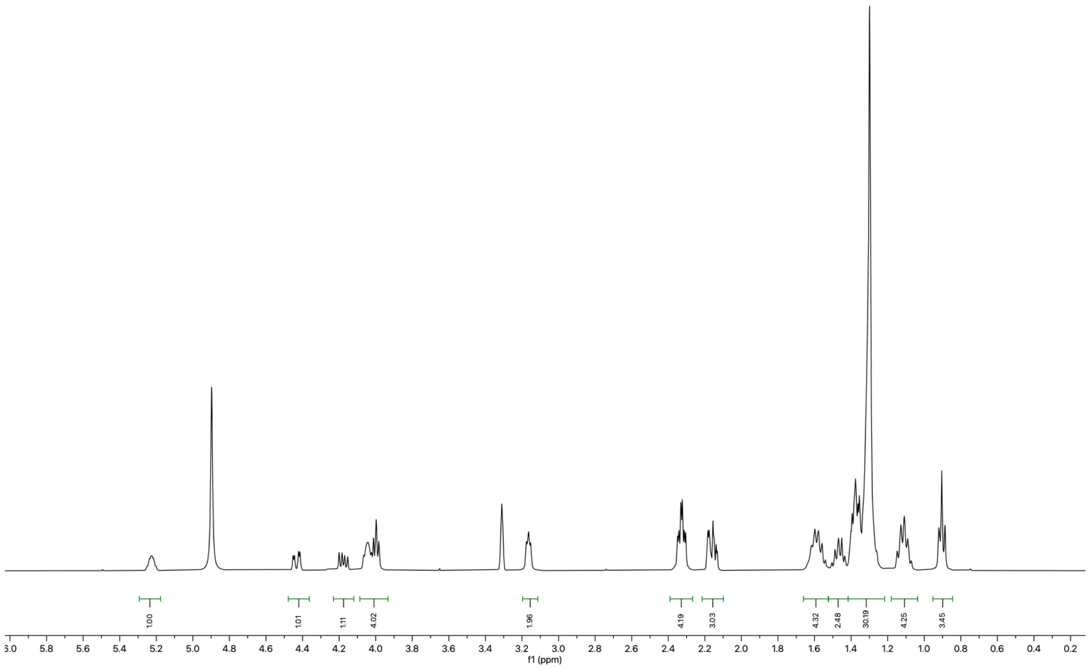

Compound **P4** $^{13}$C spectrum

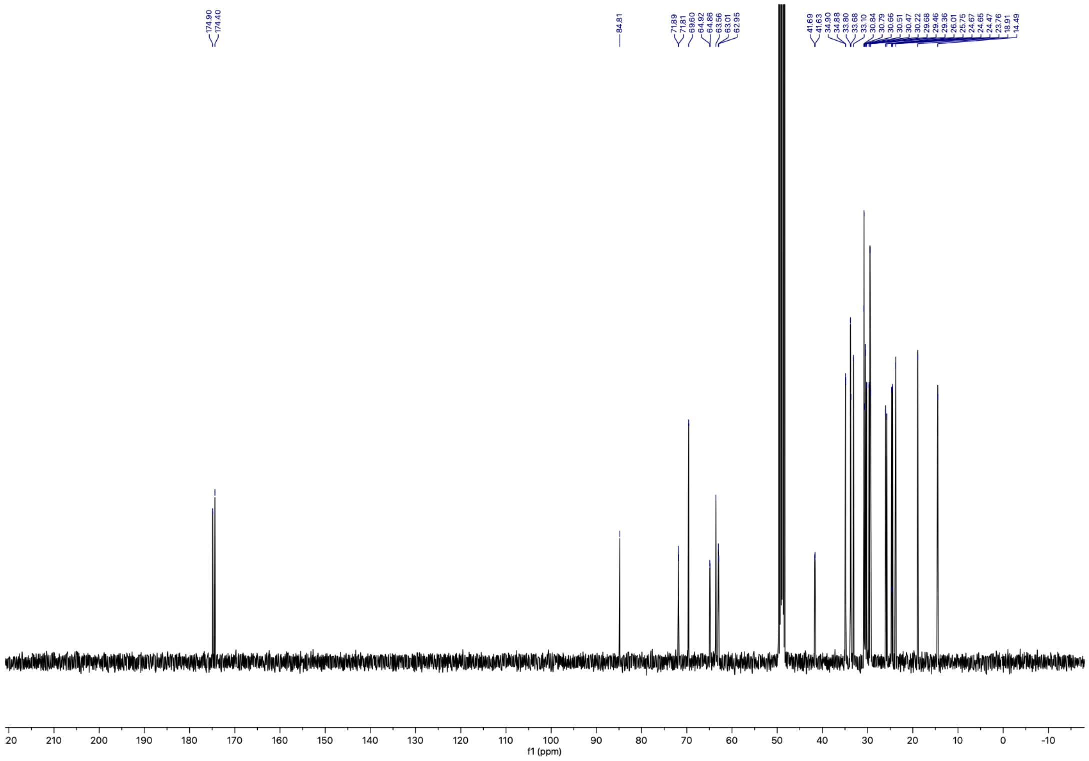


Compound **P4** $^{31}$P spectrum

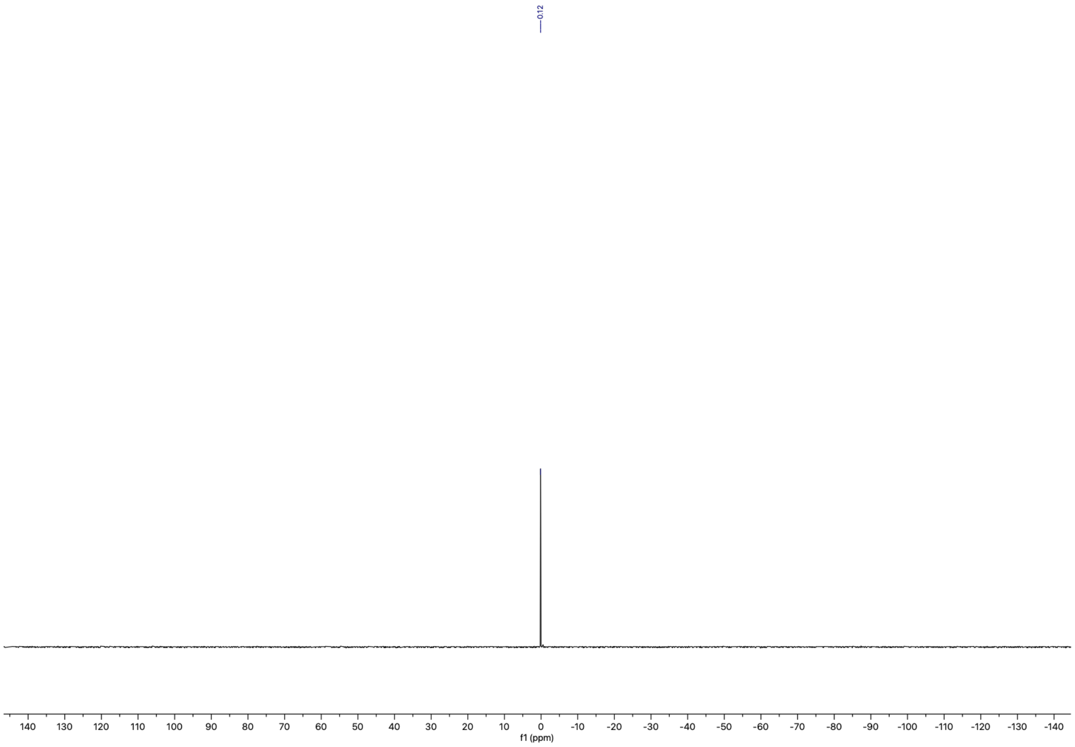
